# Mutations on a novel brain-specific isoform of PGC1α leads to extensive upregulation of neurotransmitter-related genes and sexually dimorphic motor deficits in mice

**DOI:** 10.1101/2020.09.18.300418

**Authors:** Oswaldo A. Lozoya, Fuhua Xu, Dagoberto Grenet, Tianyuan Wang, Korey D. Stevanovic, Jesse D. Cushman, Patricia Jensen, Bairon Hernandez, Gonzalo Riadi, Sheryl S. Moy, Janine H. Santos, Richard P. Woychik

**Author notes:** Co-corresponding authors: Janine Hertzog Santos, Ph.D., Richard P. Woychik, Ph.D.

## Abstract

The peroxisome proliferator-activated receptor gamma co-activator 1 alpha (PGC1α) is known as a transcriptional co-activator in peripheral tissues but its function in the brain remains poorly understood. Various brain-specific *Pgc1α* isoforms have been reported in mice and humans, including transcripts derived from a novel promoter about ∼580 Kb upstream from the reference gene. These isoforms incorporate repetitive sequences from the simple sequence repeat (SSR) and short interspersed nuclear element (SINE) classes and are predicted to give rise to proteins with distinct amino-termini. In this study, we show that a SINE-containing isoform is the predominant form of *Pgc1α* expressed in neurons. We then generated a mouse carrying a mutation within the SINE to study its functional role in the brain. By combining genomics, biochemical and behavioural approaches, we show that this mutation leads to impaired motor coordination in females, but not male mice, associated with the upregulation of hundreds of cerebellar genes. Moreover, our analysis suggests that known nuclear receptors interact with this isoform of PGC1α in the brain to carry out the female transcriptional program. These data expand our knowledge on the role of *Pgc1α* in the brain and help explain its conflicting roles in neurological disease and behavioural outcomes.

## Introduction

There is increasing interest in the role of the peroxisome proliferator-activated receptor gamma co-activator 1 alpha (PGC1α) in the brain given mounting evidence that its levels are modulated in various neurodegenerative disorders including Huntington’s (HD), Parkinson’s (PD) and Alzheimer’s disease (AD) as well as amyotrophic lateral sclerosis (ALS) (Dumont et al., 2014; Katsouri et al., 2012). However, there is only limited information about the downstream targets of PGC1α in the brain. Work done in skeletal muscle, liver, heart and brown adipos e tissue (BAT) has shown that PGC1α co-activates a series of genes prominently associated with mitochondria biogenesis, lipid metabolism, antioxidant defences and thermogenesis (Lin et al., 2004). However, conditional deletion of *Pgc1α* in the central nervous system (CNS) shows only modest changes in these processes and causes the modulation of a different set of genes associated with brain function, such as synaptotagmin 2, complexin 1 and interneuron genes (Lucas et al., 2012; Lucas et al., 2010; Lucas et al., 2014b) (Cui et al., 2006; McMeekin et al., 2018). Thus, it seems that the role and/or targets of PGC1α in the brain differ from those of peripheral tissues.

In our previous work, we identified two mouse brain isoforms of *Pgc1α* that initiated transcription from a promoter located ∼570 Kb upstream from exon 2 (Wang et al., 2016). At this position, we found a simple sequence repeat (SSR) that encoded a transcript that spliced directly to the second common coding exon of *Pgc1α* (SSR-exon2; Fig. 1A). We also identified another transcript where the SSR connected to a portion of a short interspersed nuclear element (SINE) located ∼200 Kb downstream from it, which then spliced to exon 2 (SSR-SINE-exon2; Fig. 1A). We validated the expression of these isoforms in the brain by RT-PCR. We found that the SSR-SINE-exon2 *Pgc1α* transcript was more abundant than the reference isoform in the ventral tegmental area, amygdala, hippocampus and pre-frontal cortex using publicly available RNA-seq data. Also, we found that these transcripts were brain-specific, and that the SSR and the SINE were conserved in rodents, humans, non-human primates, dogs, chickens and sticklebacks (Wang et al., 2016). Given these findings, we conclude that the SSR and SINE exons we identified correspond to the previously described human B1 and B4 exons (Soyal et al., 2012) and that the SSR-SINE-exon2 and the B1-B4 are homologous isoforms in mice and humans.

**Figure 1.**
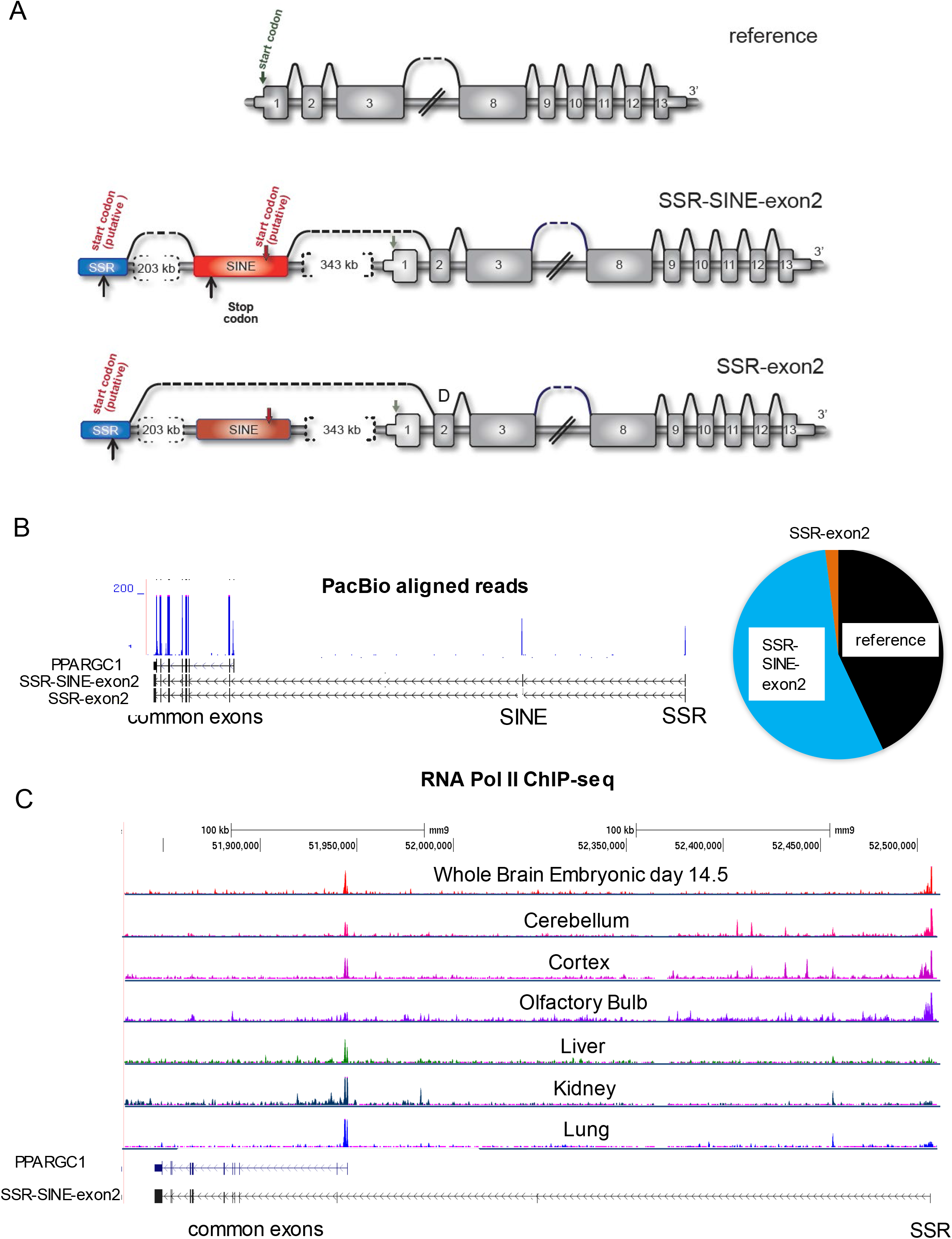
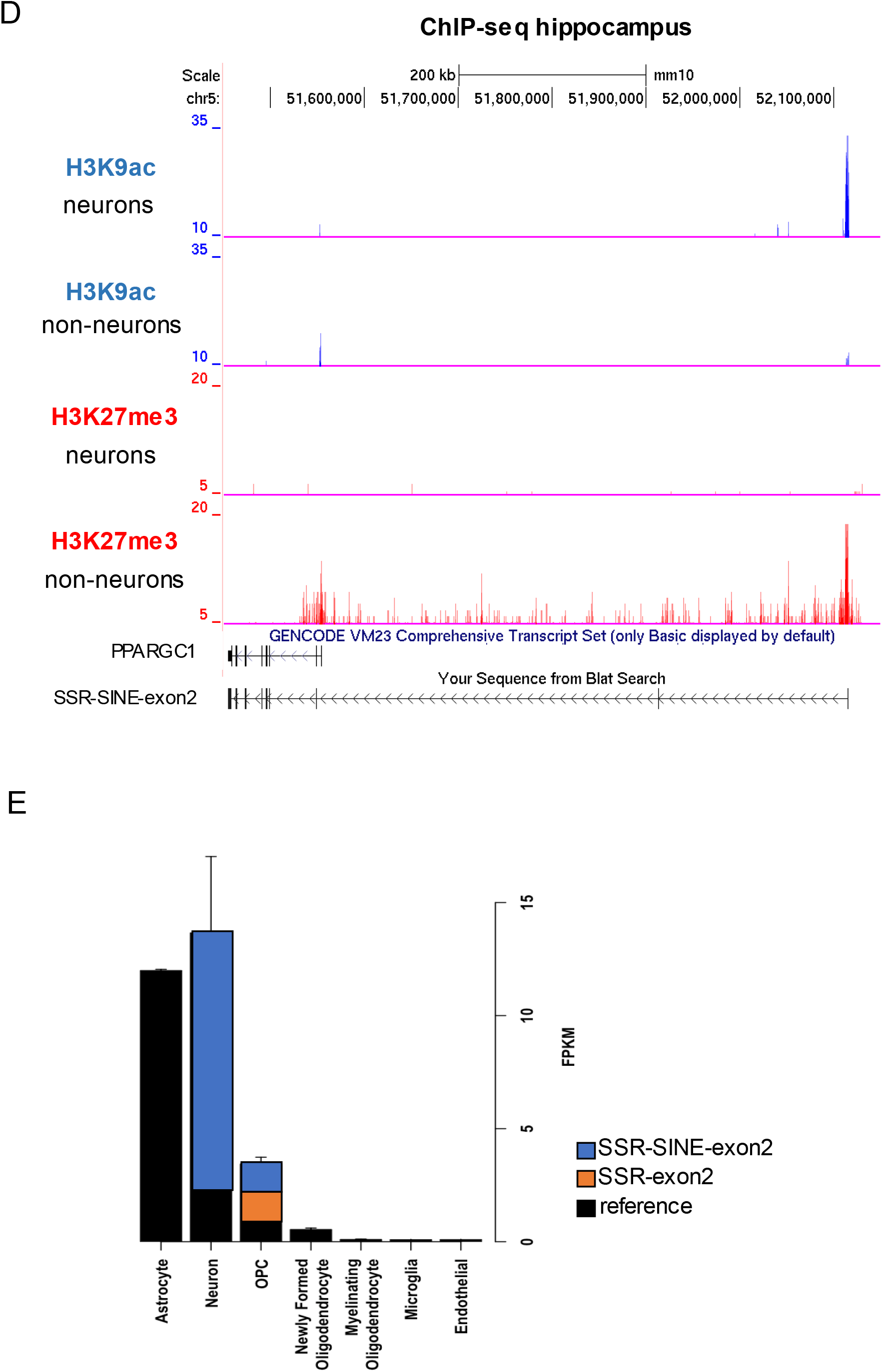
Novel isoforms of SINE-containing *Pgc1α* are expressed in the mouse brain. (A) schematic representation of gene structures of the reference, SSR-SINE-exon2 and SSR-exon2 isoforms of *Pgc1α*. Brackets depict distance between the exons, black line on top indicates splicing, grey arrow over exon 1 indicates canonical start codon. (B) UCSC browser track depicting the genomic location of SSR, SINE and exons of the reference *Pgc1α* gene (in black); gene goes from left to right. Common exons reflect those present in all isoforms, excluding exon 1 that is only present in the reference gene. In blue are PacBio peaks from reads aligning to each respective exon of the gene. Pie chart on the right depicts the proportion of full-length reads that covered the entire length of each isoform; data obtained using reads >2 Kb. (C) ChIP-seq peaks of RNA polymerase II (pol2) over the coordinates of the SSR or exon 1 of *Pgc1α* in different tissues. The genomic coordinates of the reference isoform are shown in blue on the bottom left corner. (D) Same as C but using ChIP-seq data for the promoter H3K9ac mark (in blue) or the repressive H3K27me3 mark (red) in the hippocampus. (E) Number of RNA-seq counts covering the junctions of SSR-exon2, SINE-exon2 or exon1-exon2 were used to establish the degree of expression of each of the three major isoforms of *Pgc1α* in brain-specific cell types. Data are depicted as counts per FPKM (fragments per Kb per million).

Soyal and co-workers (Soyal et al., 2012) described multiple brain *Pgc1α* transcripts that originated from an alternative promoter (referred to as B1) located ∼587 kb upstream of exon 2 in humans. They showed that the expression levels of isoforms originating from B1 were similar or higher than that of the reference gene and were confined to specific cell types. For example, while astrocytes expressed the transcript originating from the reference promoter, neurons and oligodendrocytes transcribed primarily the isoform that initiated from B1 and that contained a novel exon B4 (Soyal et al., 2012). The B1-B4 containing isoform was found to be upregulated in the striatum, cortex and in the cerebellum of mice treated with 1-methyl-4-phenyl-1,2,3, 6-tetrahydropyridine (MPTP), a drug commonly used to model PD (Torok et al., 2017). Conversely, in ALS mouse models the B1-B4 isoform seems downregulated (Bayer et al., 2017). More recently, the B1 promoter was shown to be activated by transcription factors, such as HIF1α, that do not act on the *Pgc1α* reference promoter (Soyal et al., 2020), demonstrating distinct transcriptional regulation of the brain isoforms. Finally, haplotypes encompassing the human region of the B1 promoter were associated with the age of onset of HD (Soyal et al., 2012) and with protection against PD (Soyal et al., 2019), collectively suggesting that sequence variations in brain isoforms of *Pgc1α* may contribute to disease. Nevertheless, whether they are functional and regulate similar or distinct transcriptional targets remains unclear based on these studies.

In this study, we tested the hypothesis that the SINE-containing PGC1α isoform has its own set of targets that make its function distinct from that of the reference isoform in the brain. By generating a mouse carrying a mutation on the SINE that altered the predicted ORF of the SSR-SINE-exon2 transcript, here we demonstrate that this isoform is the primary transcript of *Pgc1α* expressed in neurons and that this intragenic mutation leads to impaired motor coordination prominently in females. More importantly, we find that this mutation results in the upregulation of hundreds of genes in the female but not male cerebellum, including many involved in neurotransmission. These findings suggest that the protein expressed from the SSR-SINE-exon2, with its distinct N-terminus, functions as a sex-specific transcriptional co-repressor in the brain.

## Results

### Novel brain-specific *Pgc1α* isoforms are produced from a promoter in the SSR

The ∼600 Kb pre-mRNA of our previously identified brain isoforms of *Pgc1α* (Fig. 1A) are predicted to require between 3.3-10h to be transcribed, assuming an average transcription rate of 1-3 Kb/min (Wada et al., 2009). While it is not unusual for brain transcripts to be exceedingly large (Zylka et al., 2015), the first step in our analysis was to confirm that the full length mRNAs existed *in vivo*. We used PacBio Technology, which generates sequencing reads of up to 60 Kb in length (Rhoads and Au, 2015), to determine the types of *Pgc1α* mRNA present in the whole mouse brain. Read lengths obtained under our experimental conditions ranged from 500 bp to >5.5 Kb (Table S1). We initially analysed reads >2Kb since they would encompass the entire mRNA of the predicted novel isoforms. We found evidence for transcription of the reference, SSR-exon2 and the SSR-SINE-exon2 isoforms, with the latter being the most abundant (Fig. 1B, Table S1). No reads containing sequences upstream from the SSR were identified. When all shorter reads were analysed (see Methods), we found additional evidence for the presence of the SINE-isoform as well as other non-canonical exon-exon pairs (for details see Table S1), suggesting that there may be additional as yet uncharacterized isoforms that could be analysed.

Having confirmed that full-length transcripts occurred *in vivo*, we next determined if the SSR locus contained the promoter. To this end, we mined publicly available chromatin immunoprecipitation sequencing (ChIP-seq) data for RNA polymerase II (RNA Pol II) and for the histone H3K9ac mark, both of which are known to be characteristically enriched at promoter regions. We found RNA Pol II peaks at the SSR locus in the brain (whole brain, cortex and cerebellum) but not in the liver, kidney or lung where peaks mapped to the reference *Pgc1α* promoter (Fig. 1C). The olfactory bulb also showed an RNA Pol II peak over the SSR (Fig. 1C). Likewise, ChIP-seq data from the hippocampus showed that H3K9ac peaks were prominent over the SSR locus in neurons but not in non-neuronal cells which were enriched for the repressive H3K27me3 mark (Fig. 1D). These results show that the SSR region enriches for marks normally associated with regulation of transcriptional initiation. It is noteworthy that the SSR genomic coordinates coincides with a CpG island, which is frequently found associated with promoters in mammals, together supporting the notion that it contains the promoter of the novel *Pgc1α* brain isoforms. The H3K9ac data also suggest that the promoter at the SSR locus is primarily responsible for transcription of *Pgc1α* in neurons.

The above data prompted us to define whether expression of the different *Pgc1α* transcripts is cell type-specific in the brain. To address this, we used RNA-seq derived from distinct brain cell types (Zhang et al., 2014) and compared the abundance of reads spanning the junctions between the SSR-exon2, SINE-exon2 and exon1-exon2 to estimate the expression levels of the brain-specific transcripts with the reference *Pgc1α* isoform. Although several brain cell types were present in the dataset (Fig. 1E), only those with significant *Pgc1α* expression were considered for the analysis. We found that astrocytes expressed primarily the reference isoform since all reads from *Pgc1α* spanned the junctions between exons 1 and 2. Conversely, most reads covered the SINE-exon2 junction in neurons while in oligodendrocyte progenitor cells (OPCs) junction reads corresponding to the presence of all three isoforms were identified in similar proportions (Fig. 1E). Thus, distinct cell types express different isoforms of *Pgc1α* in the mouse brain. Most importantly, the SSR-SINE-exon2 transcript seems the primary transcript expressed in neurons.

### The SSR-SINE-exon2 isoform of Pgc1α is translated into protein

The N-terminus of PGC1α is thought to dictate its transcriptional targets (Martinez-Redondo et al., 2015; Soyal et al., 2012). Both the SSR-exon2 and SSR-SINE-exon2 isoforms skip exon 1 where the ATG used for translation initiation of the reference transcript of *Pgc1α* is present. Thus, these isoforms would need to use alternative ATGs if translated. In turn, they would give rise to proteins with different N-termini or reading frames. Analysis of the 5’ sequences of the new *Pgc1α* mouse transcripts revealed an alternative ATG within the SSR (Fig. 1A), which could connect with the ORF in the downstream exons to give rise to a protein 810 amino acids-long with 29 novel residues at its N-terminus (Fig. 2A). We predict that this same ATG would be unlikely to initiate translation of the SINE-containing transcript given a stop codon within the SINE (Fig. 1A). Downstream from this stop codon, however, is an ATG that could connect the sequences within the SINE to the ORF in the downstream exons (Fig. 1A). In this case, the resulting protein would have 6 SINE-encoded amino acids that replace the 16 amino acids at the N-terminus of the reference protein (Fig. 2A). To test whether the novel isoforms are translated into protein, we turned to publicly available ribosomal profiling data. Ribo-seq or ribosomal footprinting relies on deep sequencing of mRNA molecules after immunoprecipitation of ribosomes, giving a snapshot of the mRNAs that are actively translated within a cell (Ingolia, 2014). Thus, if these isoforms are translated into protein, the SSR and SINE, in addition to the exons of the reference protein, should be captured in the Ribo-seq dataset. We mined data derived from the hippocampus (Cho et al., 2015) and the liver (Howard et al., 2013), with the latter serving as negative control. Consistent with the repeat-containing isoforms being translated within the cell, large peaks were detected over the coordinates of the SSR, SINE and other exons from *Pgc1α* starting from exon 2 in the hippocampus (Fig. 2B). Conversely, peaks covered only exon 1 of the reference form of the gene in the liver, with no peaks over the coordinates of either the SSR or SINE (Fig. 2B, compare red and blue lines). Thus, the novel brain isoforms derived from the SSR are translated in the brain but not in the liver, as we predicted. To further confirm these findings, we then developed antibodies against epitopes unique to the amino-terminus of the predicted proteins from the SSR-exon2 and the SSR-SINE-exon2, or from the C-terminus, which would be common to all isoforms of the protein. All antibodies were highly specific to the peptides they were developed against (Fig. S1A). However, those raised against the predicted amino-terminus of the SSR-exon2 or SSR-SINE-exon2 were unspecific in tissue lysates, likely because of their short epitopes. Antibodies for the C-terminus recognized a protein of the correct molecular weight of an engineered HA-tagged PGC1α recombinant protein that we generated and expressed in NIH3T3 cells (Fig. S1B and C).

**Figure 2.**
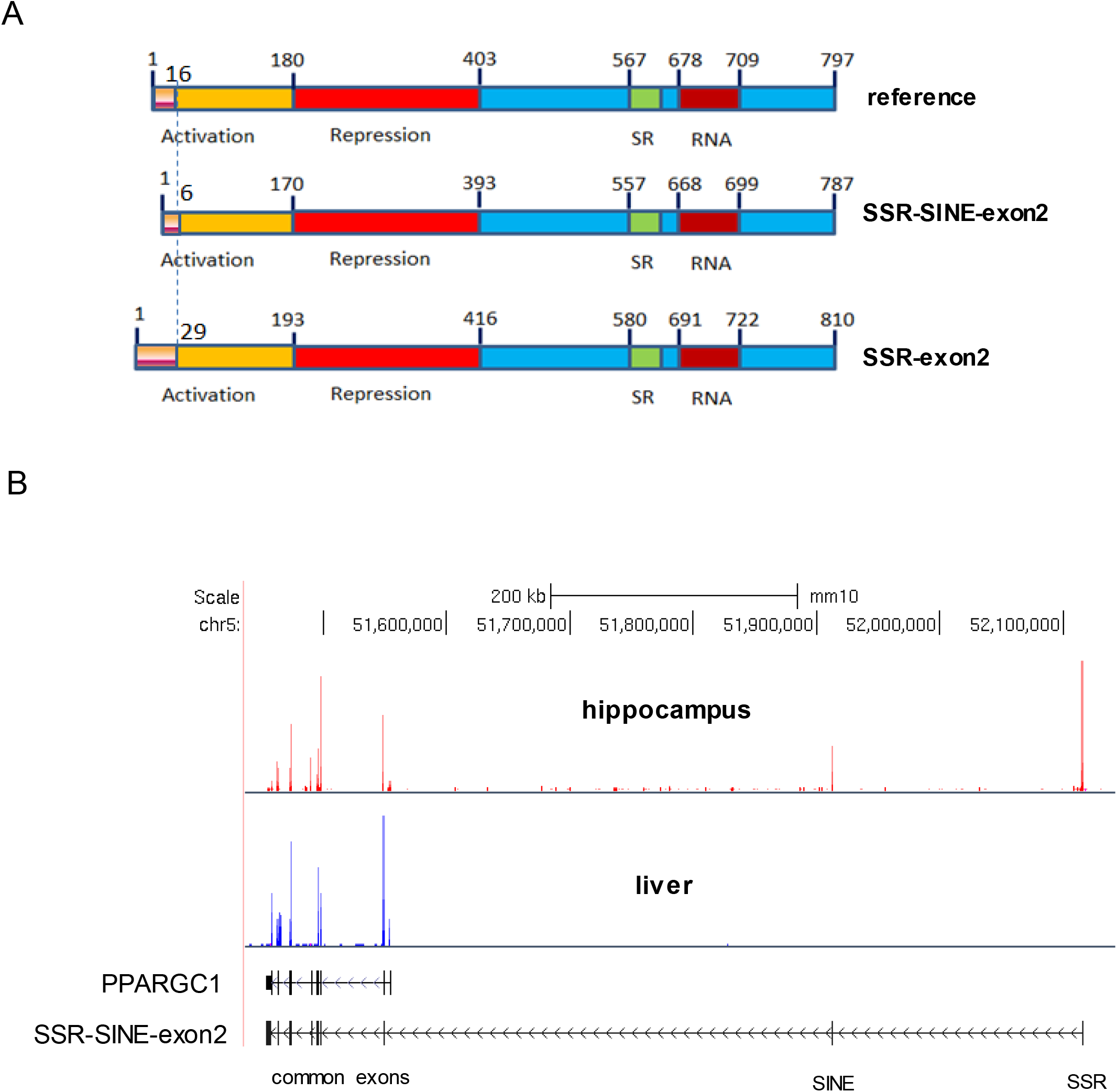
Proteins are expressed from the novel brain-specific *Pgc1α* transcripts. (A) Schematic representation of the putative protein structures of the two novel brain isoforms of *Pgc1α*; the protein derived from the reference gene is also depicted. Numbers above reflect amino acid positions; known domains are shown below. SR= serine-arginine rich. (B). Ribo-seq data from hippocampus (red) and liver (blue) were used to define the presence of the SSR, SINE and exons of reference *Pgc1α* within actively translating ribosomes. Genomic structure of the reference and SSR-SINE-exon2 isoforms are shown in black below.

### Mutation of the SINE in mice preserves normal brain anatomy but impairs behaviour and motor performance

Soyal and co-workers (Soyal et al., 2020) recently demonstrated crosstalk between the *Pgc1α* B1 and the reference promoter in the brain. Whereas they found that some stimuli activated both, which in turn seemed to co-activate each other, hypoxia was shown to engage the B1 but not the reference promoter. These results demonstrate fundamental differences in the regulation of these isoforms and suggest distinct contributions to brain physiology. However, to date there is no evidence that the new brain isoforms of *Pgc1α* are functional *in vivo*. While PGC1α KO mice that delete the common exon 3 were created (Lin et al., 2004, Leone et al., 2005, Lucas et al., 2012, 2014), these mutants eliminate all isoforms of *Pgc1α*, including the brain-specific transcripts that incorporate the SSR and SINE. Thus, to make a mouse model that could adequately establish the functional significance of the novel SSR-SINE-exon2 brain-specific isoform, we generated a mutant mouse targeting the SINE sequence. Using CRISPR/Cas9, we obtained several mutations that specifically targeted the SSR or SINE (unpublished results). We chose to establish a line of mutant mice with a 4-bp intragenic deletion immediately downstream of the putative ATG within the SINE (Fig. S2A). This mutation is predicted to abort translation of the SSR-SINE-exon2 transcript, generating a functional KO mouse. We confirmed that the transcript was still present in the brain of mutant animals and that no compensatory changes occurred in the expression of the reference isoform (Fig. S2B). Also, using the antibody generated against its C-terminus, we identified a protein in the brain of wild-type (WT) littermates but not in homozygous mutant animals with the same molecular weight as HA-tagged PGC1α (Fig. S2C). Thus, while the transcript from the 4-bp deletion mutant is present as predicted, no protein is detected in the brain of animals with this allele. We refer to these animals as SINE KO mutants.

Maintenance of this line revealed that homozygous mutant pups were born at the expected Mendelian ratio although an increase in the number of heterozygotes was noted in females (Fig. 3A). No postnatal lethality as reported with the exon 3 deletion mutants (Lin et al., 2004) was observed. Unlike for the exon 3 deletion allele, WT and SINE KO homozygotes showed no significant changes in body weight of males or females until later in life when males were ∼10% leaner (Fig. 3B). As shown in figure 3C, cresyl violet staining of sagittal brain sections did not reveal any gross anatomical abnormalities associated with alleles of the exon 3 deletion mutation (Lin et al., 2004). Likewise, we did not detect the reported spongiform lesions in the striatum (Fig. 3D) nor did we observe reduced locomotion, muscle weakness or ataxia-associated signs as described previously for the exon 3 allele (Lin et al., 2004). Thus, our data suggest that elimination specifically of the form of PGC1α expressed from the SSR-SINE-exon2 in neurons does not contribute to the post-natal lethality, neuropathology, muscle weakness and ataxia previously reported, which seem associated with the loss of the reference isoform in non-neuronal cell types. This conclusion is in line with a recent study that proposed that the neurological phenotypes are oligodendroglial in origin (Szalardy et al., 2016b).

**Figure 3.**
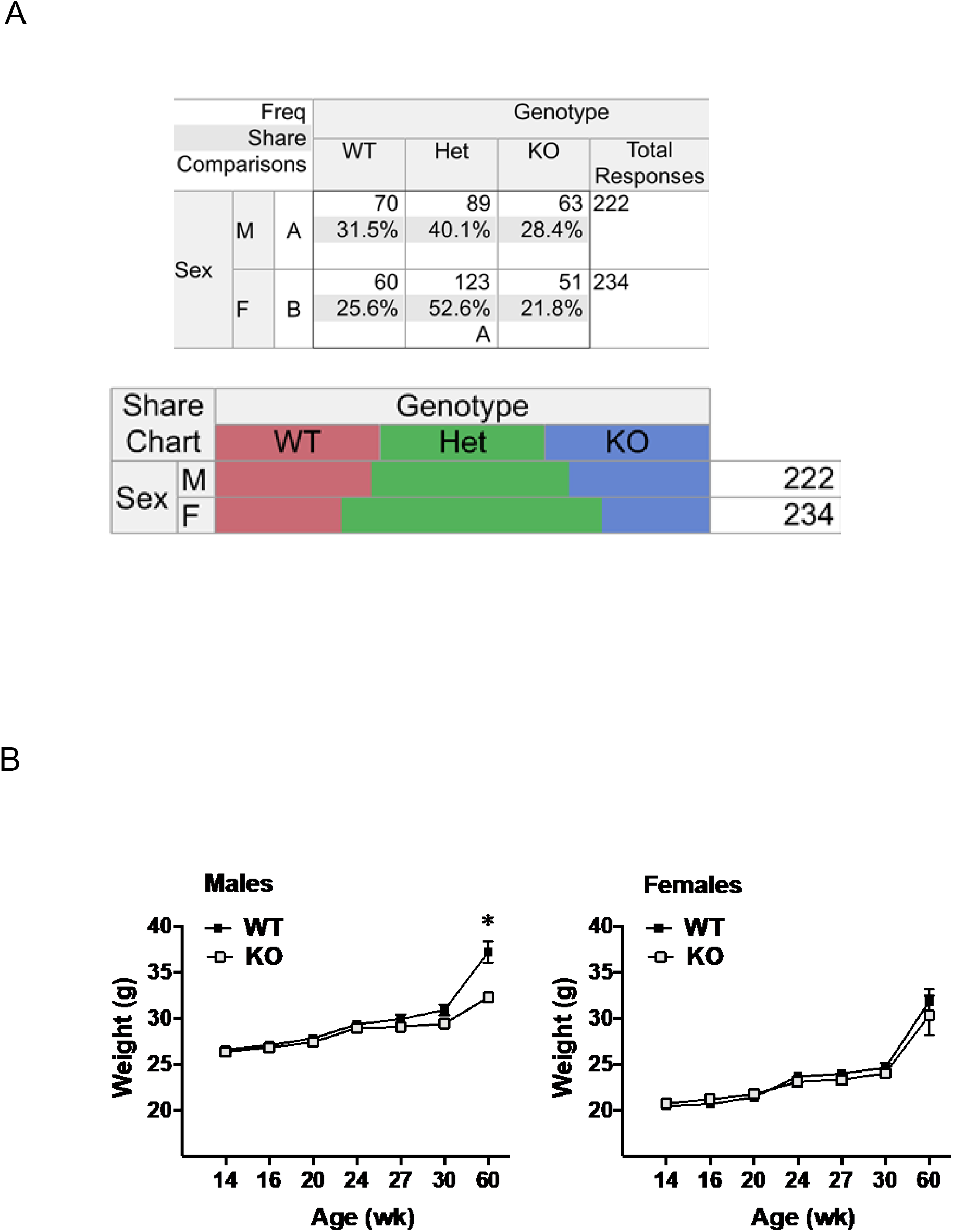
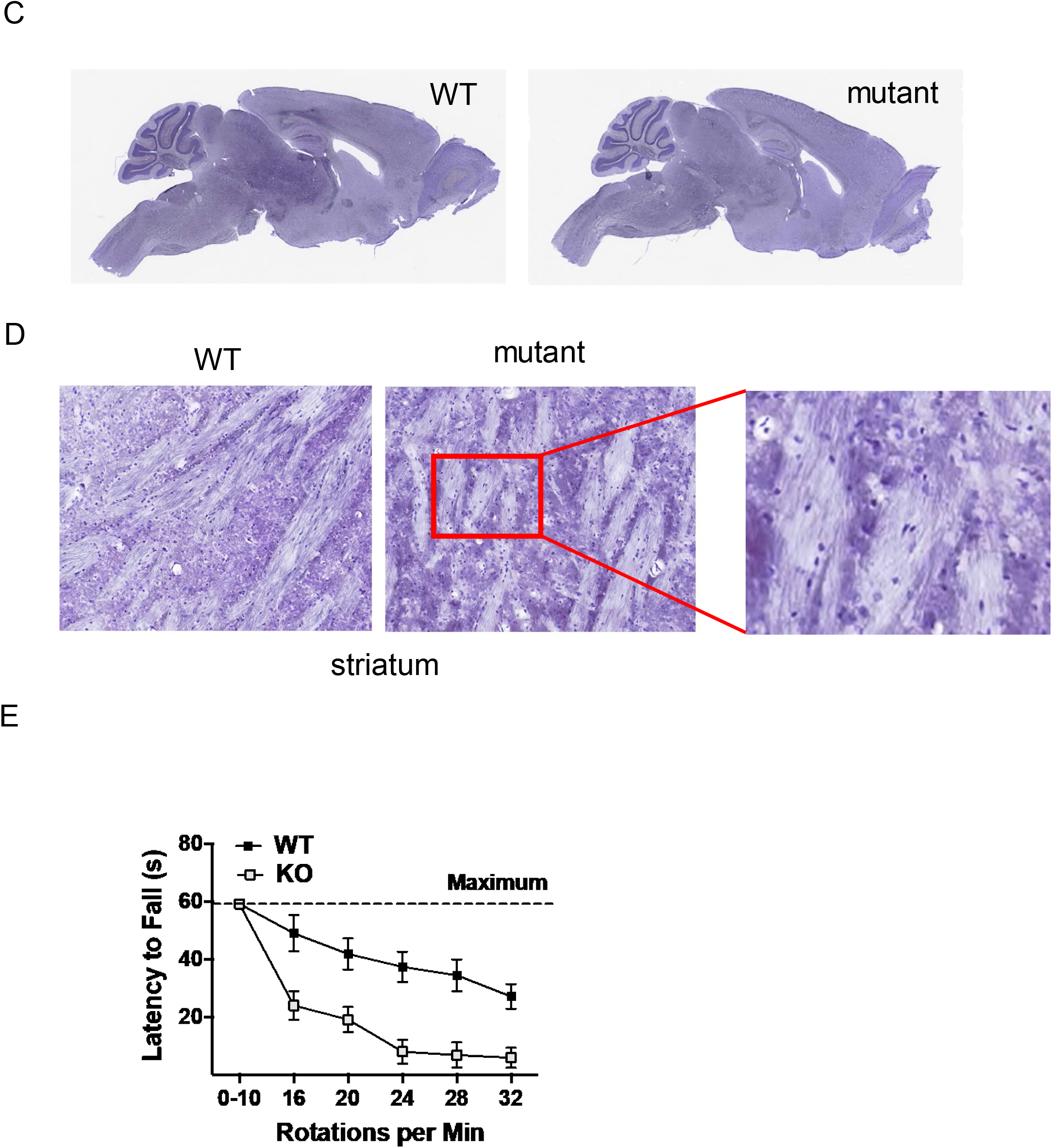
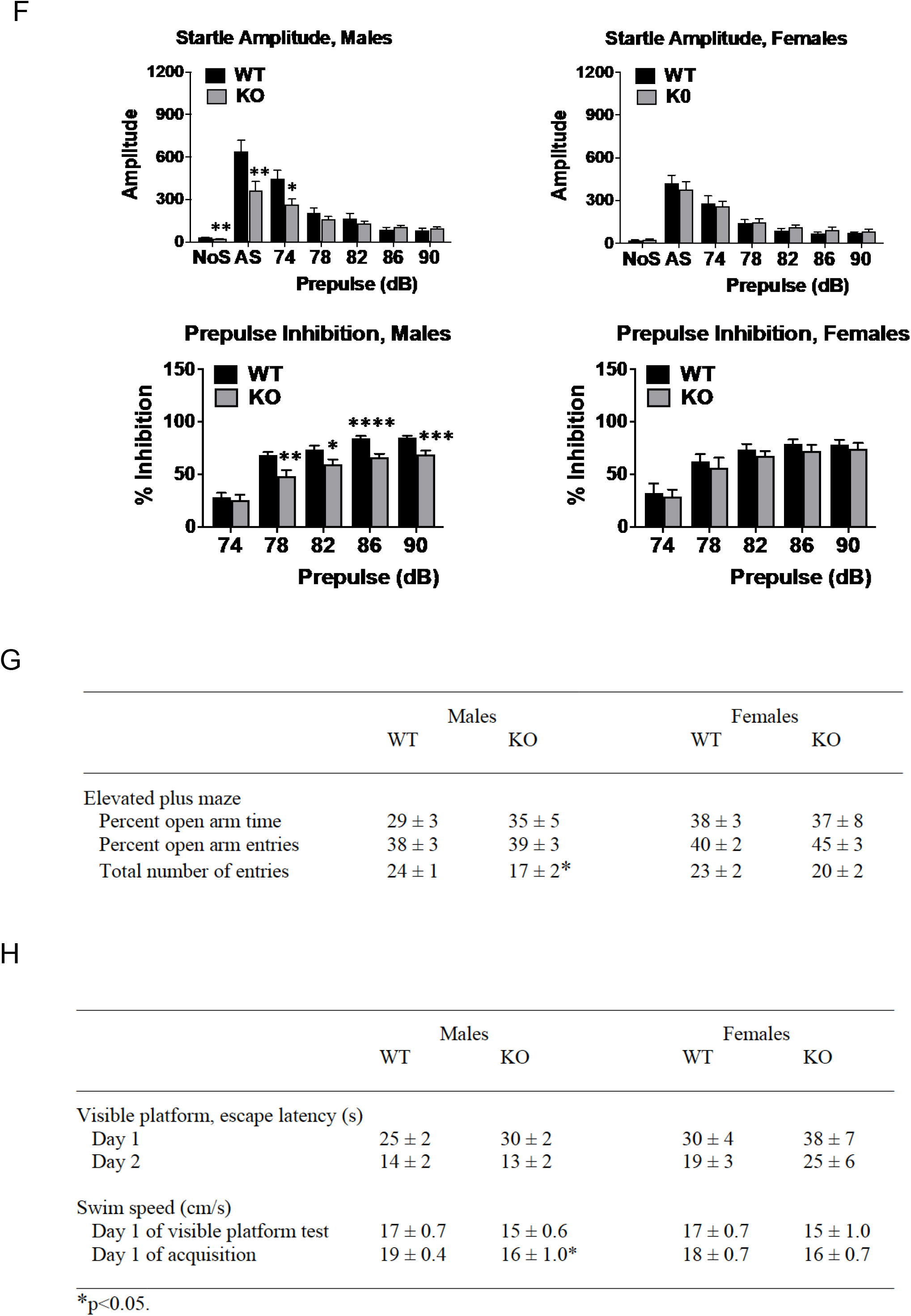
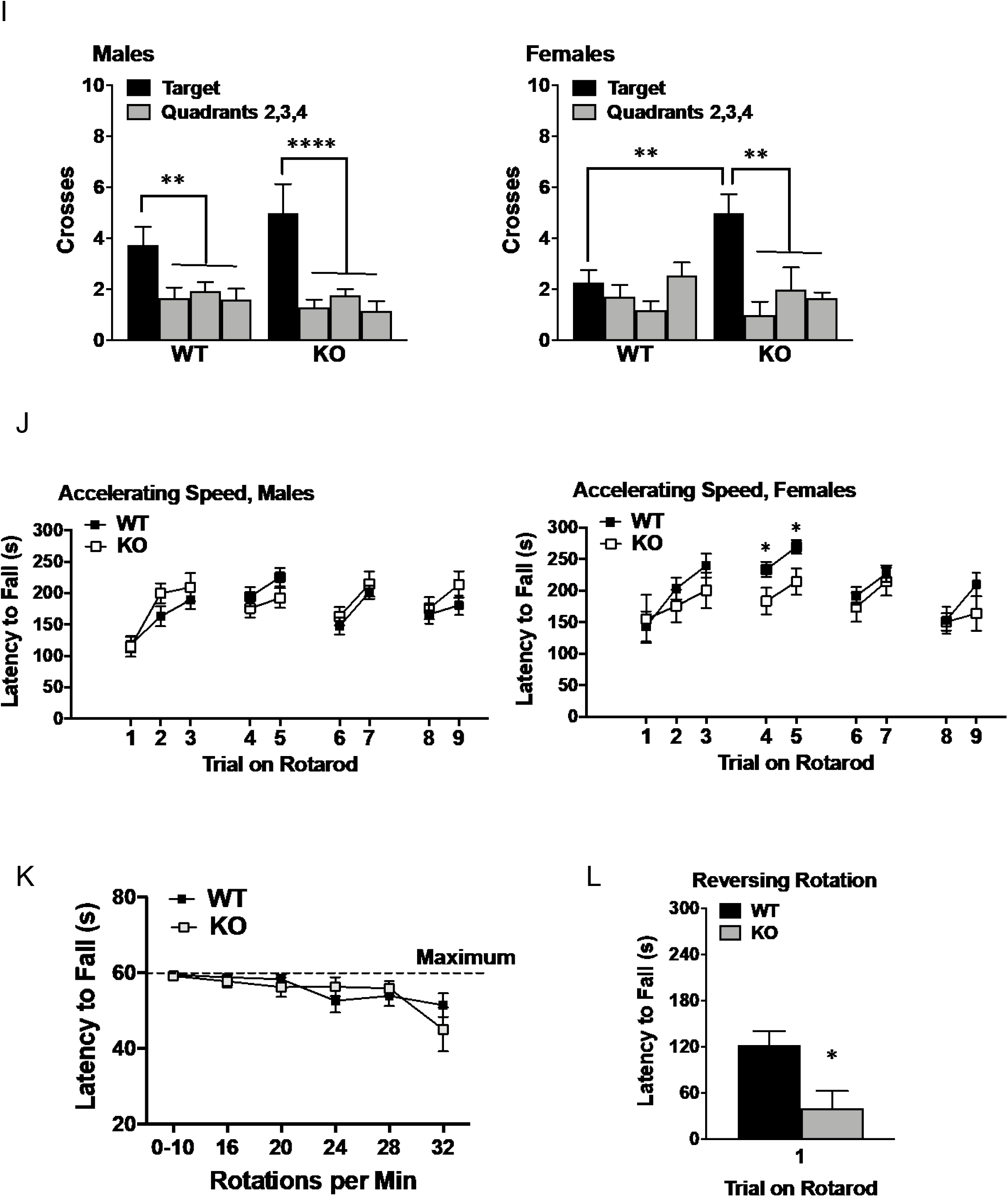
Mutations on the SINE lead to sexual dimorphic behavioural deficits in the absence of gross brain lesions. (A) Genotype frequencies by sex among alive offspring born from breeding of SINE heterozygous mutant mice, based on Pearson’s chi-square test of homogeneity (N=456 total offspring, * denotes p=0.028); WT – wild-type homozygous, Het – heterozygous KO, KO homozygous. (B) Weight changes were followed in mutant and wild-type littermates over time; final weight measure (approximately 60 weeks in age) [genotype x age interaction, F(6,156)=10.97, p<0.0001]. (C) Gross sections of brains from control or mutant littermates. (D) Rotarod test in which mice underwent 2 trials each at rotating speeds of 0–10 accelerating and 16, 20, 24, 28, and 32 fixed rpm. Each trial was a maximum of 60 sec, with at least 5 min between each trial. Data are means (+ SEM) of 2 trials per rpm. Trials 0-10 had accelerating speed; remaining trials had fixed speeds. (E) Magnitude of startle responses and (F) pre-pulse inhibition in *Pgc1α* SINE isoform-specific KO mice. Trials included no stimulus (NoS) and acoustic startle stimulus (AS, 120 dB) alone; *p<0.05. (G) Results from elevated plus maze; *p<0.05 and (H) Morris water maze as it relates to time to escape visible platform and swim speed. (I) Quadrant preference in the water was gauged after mice were given a 1-min probe trial without the platform on the final day of the hidden platform training. Measures were taken of swim path crosses over the location of the platform (in Quadrant 1), and the corresponding locations in Quadrants 2, 3, and 4. *p<0.05. (J) Motor coordination on an accelerating rotarod test as gauged in wild-type and mutant littermates. Data are means (+ SEM) for each group. Maximum trial length was 300 sec. Trials 4 and 5 were given 48 hours after the first 3 trials, when mice were 16-19 weeks in age. Mice were 31-36 weeks in age for trials 6 and 7 and were re-tested at 50-61 weeks. *p<0.05. (K) Animals were subjected to the same protocol as in (E). (L) At 56-66 weeks, animals performed a rapid reversal rotarod. Data are means (+ SEM) for each group, with males and females pooled (F(1,41) = 7.959, p = 0.007).

Behavioural changes including hyperactivity and severe impaired motor coordination in the rotarod test were associated with the brain lesions found in the exon 3 deletion alleles (Lucas et al., 2012). However, more recent studies found less severe behavioural abnormalities in brain-specific conditional alleles of the exon 3 that were deemed to be unrelated to the neuropathology (Szalardy et al., 2016a; Szalardy et al., 2018). The extent to which the SSR-SINE-exon2 isoform contributes to these phenotypes is unknown. To gain more insights into this issue, we started by subjecting SINE KO homozygotes and their WT littermates to the same rotarod assay applied by Lucas et al., 2012, which involves two trials per day of increasing rotational speeds from 16-32 RPM. SINE mutants performed very poorly in this test (Fig. 3E), essentially phenocopying the defects previously described with the exon 3 deletion (Lucas et al., 2012). These results suggest that loss of the SSR-SINE-exon2 *Pgc1α*, which is the primary isoform expressed in neurons, functionally contributes to decreased rotarod performance in a way that is unrelated to the presence of lesions in the brain. We then subjected an independent cohort of animals to a battery of behavioural tests, including a more conventional and less challenging rotarod (see below). Table 1 summarizes the timeline and overall test results obtained. While no statistical differences were found between WT and SINE KO homozygous mutant animals for most protocols (Fig. S3), a few sexual dimorphic outcomes were identified in the mutant animals. For example, male but not female SINE KO homozygotes had significant decreases in the magnitude of the startle response and impaired pre-pulse inhibition (Fig. 3F). Pre-pulse inhibition is disrupted in several neuropsychiatric disorders, including schizophrenia, which not only has a male preponderance but also has been associated with reduced cortical expression of PGC1α (McMeekin et al., 2016). Male SINE KO homozygotes also showed decreased arm entries in the elevated plus maze (Fig. 3G) and slightly slower initial swim speeds in the water maze (Fig. 3H), which may be indicative of increased anxiety and impaired swimming-related motor coordination, respectively. Interestingly, conflicting findings have been reported on anxiety-like behaviours for the full body exon 3 deletion alleles (Leone et al., 2005; Szalardy et al., 2018). Finally, we found that female, but not male SINE KO homozygous mutants, had significantly higher target quadrant preferences relative to control animals in the Morris water maze (Fig. 3I), suggesting improved spatial learning and memory (Vorhees and Williams, 2014).

**Table 1.** Behavioral testing regimen.

| Table 1 |  |  |
| --- | --- | --- |
| Behavioral testing regimen |  |  |
| Age (week) | Procedure | Outcome |
| 14-17 | Elevated plus maze test for anxiety-like behavior | Males - Reduced overall entries |
| 15-18 | Locomotor activity and exploration in a 1-hour open field test | No differences |
|  | Wire-hang test for grip-strength | No differences |
| 16-19 | Accelerating rotarod test | Female – impairments in trials 4 and 5 |
|  | Trials 1-3 (first test); trials 4 and 5 (second test, 48 hr later) |  |
| 17-20 | Social approach in a three-chamber choice task | No differences |
| 18-21 | Marble-bury assay for anxiety and perseverative responses | No differences |
|  | Prepulse inhibition of acoustic startle responses | Males - Reduced startle and PPI |
| 19-24 | Buried food test for olfactory ability | No differences |
| 20-25 | Morris water maze; visible platform test | Males - Reduced swim speed |
| 21-27 | Morris water maze; acquisition of spatial learning | Females – improved probe trial |
| 23-28 | Second acoustic startle test | No differences |
|  | Hot-plate test for thermal sensitivity | No differences |
| 26-31 | Conditioned fear test for contextual and cue learning | No differences |
| 28-33 | Second fear test for memory retention | No differences |
| 31-36 | Accelerating rotarod, trials 6 and 7 | No differences |
| 32-37 | Accelerating and fixed speed rotarod (1 min trials) | No differences |
| 50-61 | Accelerating rotarod, trials 8 and 9 | No differences |
| 56-66 | Rapid-reversal rotarod test (2 trials) | Males and females - impaired |
| 58-67 | Fixed speed rotarod (2 min trials) | No differences |

At 16-19 weeks in age, WT and SINE KO homozygous littermates were subjected to a standard accelerating rotarod protocol that differed from the test that we employed initially (Fig. 3E). This protocol consisted of sessions over multiple days in which speed progressively increased from 3 to a maximum of 30 rpm across 5 min. The first test consisted of 3 trials of 5 min with 45 sec in between each (Fig. 3J, 1-3), which were followed by two additional trials that occurred 48h later (Fig. 3J, 4-5). Females but not males showed significant impairments at trials 4-5 (Fig. 3J). At weeks 31-36 (trials 6-7) and 50-61 (trials 8-9), these animals were retested but no differences between WT and mutant littermates were observed (trials 6-7 and 8-9, Fig. 3J). These results show that the motor learning deficit in this less difficult assay was female-specific and more subtle than in the previously reported protocol that we initially used (Lucas et al., 2012).

This argues that females may be more sensitive than males to the loss of the SSR-SINE-exon2 isoform. In addition, the absence of the motor deficit in the later tests (trials 6-7 and 8-9) suggests that motor learning, and perhaps previous experience in other assays such as the water maze (see Table 1 for timeline), allows them to overcome this deficit.

To test the sex-dependence of the rotarod deficit and the potential role of motor learning in overcoming it, we took two different approaches. Firstly, we subjected these same animals to the more challenging rotarod test based on Lucas et al., 2012 (as in Fig. 3E). Consistent with the hypothesis that prior motor learning allowed them to overcome the deficits on this task, no impairments were identified (Fig. 3K). We then subjected them to a difficult protocol to which they had not been exposed, which involved rapid reversals in the direction of rotation of the rotarod. This revealed deficits in both males and female mutants (Fig. 3L). They were then re-tested on an easier rotarod version with fixed speeds and no differences were observed (see Table 1). Given that animals were tested at about the same ages in the latter three tests (Table 1), the deficit in the rapid reversal task is likely to be driven by task difficulty and novelty rather than an age-related decline. Taken together, these data indicate that deficits in motor coordination and motor learning can be unmasked in the SINE KO mutants when animals are exposed to a novel and/or difficult protocol, that females are more affected and that training experience can overcome these deficits.

### SINE-mutant female mice exhibit increased neurotransmitter-associated gene expression in the cerebellum

The phenotypes observed in the rotarod tests suggest a cerebellar but not a striatal-dependent deficit since the latter is more important for motor learning (Dang et al., 2006). Although prior studies reported cerebellar alterations in the exon 3 deletion mutant animals (Lucas et al., 2014b), the SINE KO mutants did not show obvious brain anatomical defects (Fig. 3C, D). To better understand the molecular underpinnings of the rotarod phenotype, we next profiled gene expression in dissected cerebella (Cb) of age-matched WT and SINE KO homozygous littermates, as well as the rest of the brain (Br, whole brain minus cerebellum) by microarray assays. We analysed differential expression based on RMA-normalized probe intensities by the LSTNR method (Lozoya et al., 2018) (2^3^ full-factorial design, N=2 per group). Principal component analysis of 12,527 multivariate significant probe sets showed that the largest statistical differences between group means were observed in the cerebellum, most notably in females (Fig. 4A). Indeed, less than 50 probes were differentially enriched in the rest of the brain for each sex (Fig. 4B lower panels), whereas 2,016 probes in females or 354 probes in males were different between cerebellum of WT and SINE KO mutant littermates (Fig. 4B upper panels). Most probes showing significant expression differences in female cerebella were predominantly upregulated in the mutants (Fig. 4B, upper left panel). In males, the probes that were different between WT and mutant animals were mainly downregulated (Fig. 4B, upper right panel), which is consistent with what is known about loss of the reference PGC1α isoform in peripheral tissues. These data point to sex differences associated with the loss of the protein derived from SSR-SINE-exon2 transcript and, most importantly, suggest that this isoform normally represses gene expression in the female brain. The degree of probe overlap between samples can be found in Fig. S4; the list of the 1,615 genes encompassed by these probes can be found in Table S2.

**Figure 4.**
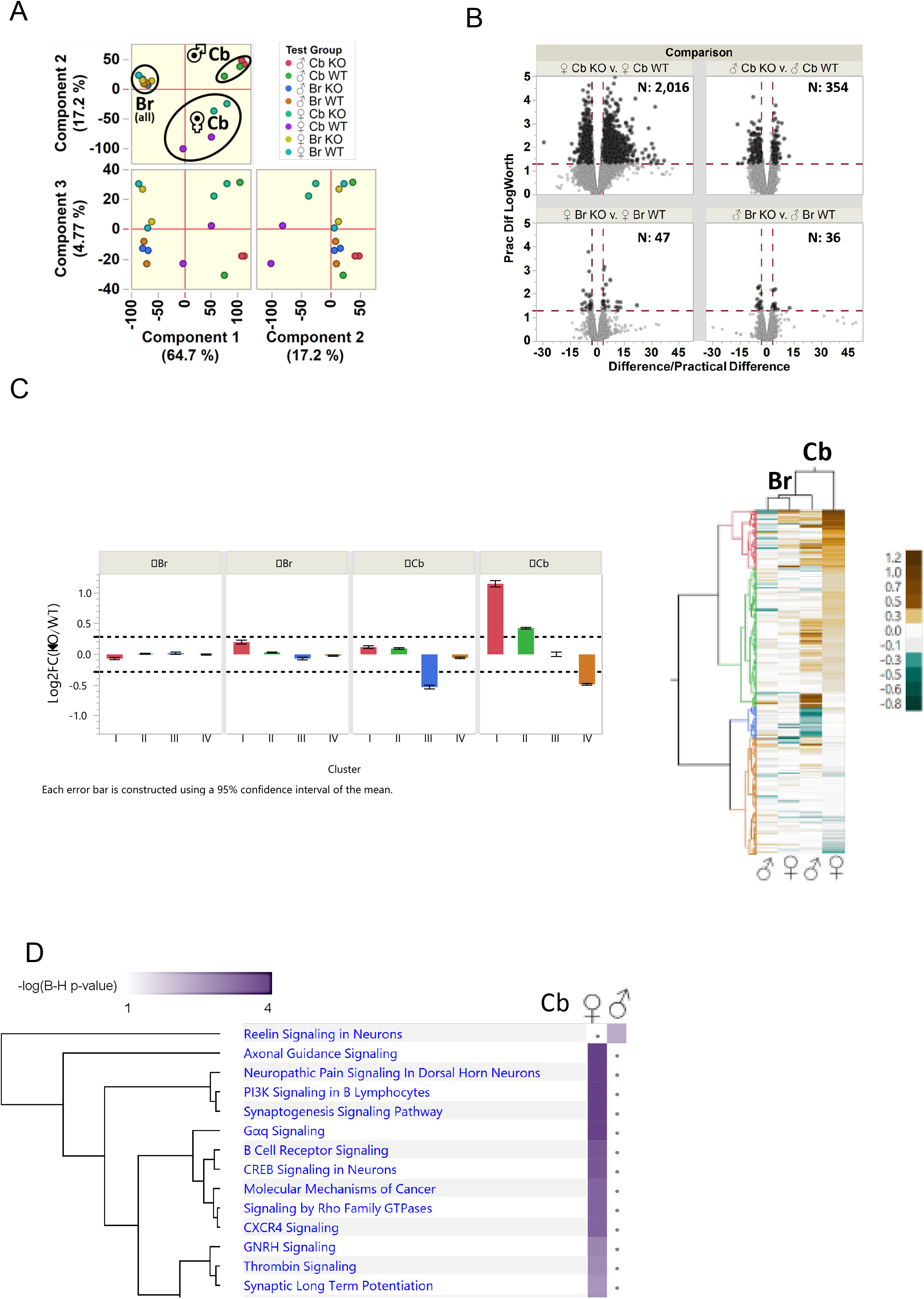
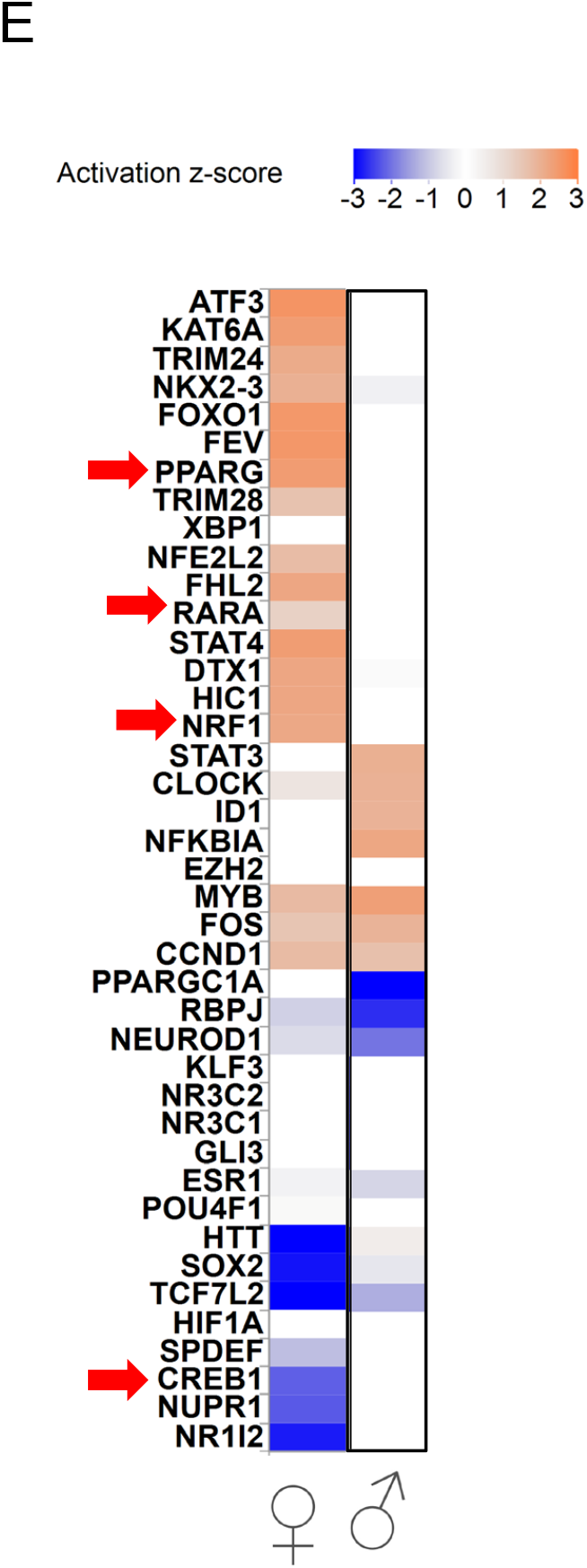
Loss of the SINE isoform leads to sexual dimorphic gene expression profiles. (A) Principal component analysis of 12,527 multivariate significant probe sets (RMA-normalized probe intensities, log_2_FC ANOVA FDR p<0.05, |log_2_FC|>0.286 for SNR>1 vs. grand mean; PC1_organ_ + PC2_sex_ = 82% total variance) was used to determine the main components driving the differences between gene expression profiles of wild-type and mutant littermates. (B) Volcano plots depicting 9,996 statistically differential probe sets between any genotype × organ × sex groups overall (multivariate significant, log_2_FC *post hoc* pairwise p<0.05; 5% practical difference |log_2_FC|>0.094); black highlights 2,363 differential probe sets combined that meet differential criteria in same-organ, same-sex comparisons between WT vs. SINE KO mutant mice, the number of which is indicated inside each panel. (C) Gene expression data was segregated into patterns using unsupervised hierarchical clustering based on 1,980 single-gene probe sets encompassing 1,615 gene annotations, corresponding to the subset from same-sex 2,363 differentially expressed probes curated against RIKEN cDNA clones, multi-gene, or no-gene annotations. The heatmap on the right (Cb = cerebellum, Br = rest of brain, i.e. whole brain minus cerebellum) depicts the patterns giving rise to the distinct clusters: red cluster I (330 single-gene probe sets, 257 genes, log_2_FC min: +0.12, max: +3.48, IQR: [+0.84, +1.31]), green cluster II (807 single-gene probe sets, 677 genes, log_2_FC min: –0.32, max: +0.92, IQR: [+0.34, +0.55]), blue cluster III (172 single-gene probe sets, 156 genes, log_2_FC min: –1.53, max: –0.10, IQR: [–0.63, –0.38]), and orange cluster IV (671 single-gene probe sets, 591 genes, log_2_FC min: –2.79, max: +0.27, IQR: [–0.57, –0.37]); cluster-wise expression differences were deemed statistically robust based on |log_2_FC|>0.286 (SNR>1 threshold). (D) Ingenuity Pathway Analysis (IPA) of based on the curated list of 1,980 single-gene differential probe sets with statistically robust expression differences per group (|log_2_FC|>0.286; male cerebellum: 457 probe sets; female cerebellum: 1,738); dots indicate pathways without significant enrichment within gene sets per group. (E) Upstream regulators predicted based on the genes differentially expressed in females (left) or males (right). All differentially expressed genes, only the ones with a PGC1α recognition sequence within ± 1Kb of the annotate promoter or the ones without it were separately considered. Only data from cerebellum was analysed. Z-scores range depicting activation (orange) or inhibition (blue) are shown.

We also submitted the differentially expressed genes to unsupervised hierarchical clustering to define patterns of expression. This approach revealed 4 different dominant patterns that were binned into clusters I through IV (Fig. 4C). Again, the largest differences in the degree of gene expression changes were observed in female cerebella, involving genes within clusters I and II that were all upregulated (257 and 677, respectively, Fig. 4C). These genes involved many processes specific to the brain, including receptors, transporters and biosynthetic enzymes of the neurotransmitters glutamate, dopamine, serotonin and cholinergic synapses (Table S2). The second largest changes occurred in genes binned into clusters III in males (156) and IV in females (591), which were downregulated (Fig. 4C). These were involved in different cellular functions and included kinases, phospholipases, some transporters and channels in addition to immune-associated genes. It was noteworthy that no mitochondrial, antioxidant or other genes previously identified as differentially expressed in the cerebellum of animals with the conditional exon 3 deletion allele (Lucas et al., 2014b) were identified (Table S2). These may reflect the loss of all isoforms of *Pgc1* in the brain, supporting the notion that the proteins derived from the SSR-SINE-exon2 and from the reference *Pgc1* transcripts are not functionally equivalent in this tissue. To more broadly understand the impact of loss of the protein expressed from the SSR-SINE-exon2 isoform, we used Ingenuity Pathway Analysis (IPA) to identify biological processes enriched based on the cerebellar gene expression profiles. Given the large difference in the number of genes and their sex-specific up or downregulation, we performed IPA separately in males and females. Several pathways relevant to brain physiology were enriched in females, while only reelin signalling was identified in males (Fig. 4D). However, it is interesting to note that the first *reeler* KO mouse described showed severe cerebellar abnormalities (Caviness, 1976). Next, we used KEGG (Kyoto Encyclopedia of Genes and Genomes) to identify pathways enriched based on upregulated versus downregulated genes. Based on upregulated genes, we found over 50 significantly represented KEGG pathways, many of them relevant to brain-specific processes including the first top 7 (Table S3). Notably, the top category involved glutamatergic synapses, consistent with glutamate being one of the most abundant neurotransmitter in cerebellar cells (Zampini et al., 2016). Increased glutamate signalling was recently reported in the neocortex and hippocampus of another conditional brain-specific PGC1α mutant that involved the deletion of the common exon 3 (McMeekin et al., 2020). In contrast, only 3 pathways were significantly enriched based on downregulated genes, none of which were unique to brain physiology (Table S3).

The reference PGC1α isoform is known for interacting with a set of nuclear receptors (NRs) to regulate downstream targets in peripheral tissues, including the estrogen receptor α (ERα). Given the sexual dimorphism identified in the gene expression profiles, we started by asking whether estrogen through the ER could be involved in this response. Using HOMER, a motif analysis algorithm, and a window of ±1 Kb from the transcriptional start site (TSS) of genes, we found that about 18% of differentially expressed genes (DEGs) had an estrogen responsive element (ERE, Table S2). KEGG analysis of these 290 genes demonstrated that they enriched for 4 brain-specific pathways, although glutamatergic synapse was not among them (Table S4). To identify other potential proteins that could function as co-regulators of the gene expression with the protein expressed from the SSR-SINE-exon2 isoform, we used IPA to predict the upstream drivers of the transcriptional program. IPA derives its prediction from established interactions between transcription factors (TFs) and target genes based on published experimental evidence. Using exclusively the female gene expression, we identified many TFs that were previously associated with stress response or inflammatory signalling (Fig. 4E). It was noteworthy to find NRs known to interact with the reference PGC1α such PPARG (peroxisom e proliferator receptor gamma), NRF1 (nuclear respiratory factor 1), the RARα (retinoic acid receptor alpha) and CREB1 (cyclic AMP responsive element binding protein 1) (Fig. 4E). These NRs were unexpected because their classic downstream targets were not present within our dataset. Nevertheless, it is possible that these NRs regulate target genes in the brain that differ from those in peripheral tissues. Consistent with this possibility, PPARG has been shown to modulate NF-Kβ immune-dependent gene expression in microglia (Bernardo and Minghetti, 2006), an effect that is not observed in the periphery.

To further explore the role of NRs specifically in the female cerebellum, we employed HOMER to identify the extent to which the identified DEGs harboured response elements (RE) that could be recognized by those NRs. We found that 20% of genes had a RE for NRF1, 13% for CREB1 and 2% for RARα (Table S2), but when using these genes, no pathway enriched with a significant adjusted p-value (Table S4). About 56% of DEGs had a PPARG binding site (Table S2), 457 of which were not only upregulated but enriched for the same brain-specific pathways as the 884 upregulated genes; glutamatergic synapse was the top pathway (compare Tables S3 and S4). Statistical analysis revealed that such a high enrichment for PPARG recognition sequence was not significantly different than the abundance of these sites in the other 13,509 transcribed but not differentially expressed cerebellar genes (Table S2). Despite all of this, about 100 genes that we identified as having a PPARG response element using HOMER are predicted to be downstream targets based on simulations on the PPAR database (Fang et al., 2016). It is also noteworthy that PPARG has been shown to drive sexual dimorphic phenotypes in the periphery and in the brain (Duan et al., 2010; Park and Choi, 2017), including in models in which PPARG agonists were employed (Benz et al., 2012). Taken together, these data support the hypothesis that a component of the sexual dimorphic gene expression program identified in the SINE KO mutant cerebellum may be related to disruptions in the interaction between the SINE-containing protein and some of the same NRs that PGC1α interacts with in peripheral tissues.

## Discussion

Our understanding of the function of PGC1α in the brain is still emerging. Despite findings describing multiple and uniquely regulated transcriptional PGC1α isoforms in the brain (Soyal et al., 2020; Soyal et al., 2012; Soyal et al., 2019), their function remain largely unknown. Full body and CNS-conditional *Pgc1α* KO mice were generated over a decade ago and found to have neurological phenotypes, including altered behaviour, but inconsistent data has been reported by different groups (Dougherty et al., 2014; Leone et al., 2005; Lin et al., 2004; Lucas et al., 2012; Lucas et al., 2010; Lucas et al., 2014b; McMeekin et al., 2018). Because those KO strains were generated by deletion of exon 3, which is common to all transcripts, it was impossible to dissect the potential contribution of the different isoforms to these phenotypes. In this study, by characterizing animals devoid solely of the novel SSR-SINE-exon2-encoded protein, we not only identified that this isoform inhibits genes but also that it drives a sex-dependent brain transcriptional program. These findings strongly suggest that the different brain *Pgc1α* isoforms are not functionally equivalent, which may further help explain conflicting reports about the benefits or detriment of modulating levels of the canonical PGC1α in the context of neurodegenerative disease (Ciron et al., 2012; Clark et al., 2012). Most notably, our data suggests that the protein expressed from the SSR-SINE-exon2 isoform functions as a transcriptional repressor while the protein from the reference isoform functions as a transcriptional co-activator.

One of the best examples of the differential function of the *Pgc1α* brain isoforms stems from our findings that the SINE mutant mice did not show the gross neuroanatomical changes previously reported for the exon 3 deletion mutants. Yet, loss of this specific isoform resulted in significantly altered gene expression in the female cerebellum consisting primarily in the upregulation of genes, including those associated with neurotransmission. Furthermore, this in line with greater cerebellar dysfunction in the female SINE KO mice relative to males, as evidenced by notable motor coordination defects in the less difficult rotarod protocol (Mason and Sotelo, 1997). This sexual dimorphic phenotype is reminiscent of the motor impairments reported in models of accelerated aging and PD (Antzoulatos et al., 2010; Baeta-Corral et al., 2018). Nevertheless, the underlying cause for the sexual dimorphism observed herein remains unclear. Our analysis identified that 20% of the DEGs are potential targets of ERα; PPARG may be involved in the regulation of another ∼20-60% of them. Interestingly, 62% of the genes containing an ERE in our dataset also bear a PPARG recognition sequence (Table S2), suggesting that some ERα target genes may be regulated by PPARG. This is in line with previous reports demonstrating that PPARG can mediate the expression of estrogen target genes (Keller et al., 1995; Nunez et al., 1997). Recent data demonstrated that PPARG is highly expressed in neurons in the adult mouse brain (Warden et al., 2016), but little is still known about its downstream targets in this cell type. In fact, the effects of PPARG on brain physiology, other than neuroinflammation, are poorly understood and have been mostly inferred using agonists such as pioglitazone, which has been shown to improve neurological deficits in different disorders, including rotarod performance in a mouse model of AD (Toba et al., 2016). Additional experiments are required to test the crosstalk between PPARG, the ERα or estrogen and the protein expressed from the SSR-SINE-exon2 isoform of *PGC1α* in the brain.

The comprehensive upregulation of a cerebellar gene expression program by the loss of the protein expressed from the SSR-SINE-exon2 has not been reported previously using the exon 3 deletion of PGC1α. It is possible that the upregulation of genes in the absence of PGC1α in the brain was missed because most studies used RT-PCR to identify differential gene expression, which by its nature would limit the number and types of genes interrogated (Lucas et al., 2012; Lucas et al., 2010; Lucas et al., 2014b; McMeekin et al., 2020). Also, the use of the exon 3 mutant, by virtue of ablating all isoforms of PGC1α in the brain, may have masked this phenotype. Alternatively, but not mutually exclusive, historical bias may have skewed previous results by emphasizing the analysis of repressed genes. Interestingly, two recent studies showed that the loss of PGC1α in the brain led to the upregulation of genes, including in the striatum and in the hippocampus. In the striatum, upregulation of 429 genes was noted but only the 659 downregulated genes were further studied (McMeekin et al., 2018). Our re-analysis of the striatum data using different statistical criteria showed that over 1,000 were upregulated and they were enriched for some of the same brain-relevant pathways as identified by us; interestingly, about 50% of them had a PPARG recognition element (Table S5). Thus, it seems that expression of the SSR-SINE-exon2 containing isoform of PGC1α is tied to normally repressed gene expression profiles in the brain.

Glutamatergic synapse was the top category enriched by the upregulated genes (Table S3). A recent study using the exon 3 deletion mutant found ambulatory hyperactivity in response to a novel environment and enhanced glutamatergic transmission in the neocortex and hippocampus, along with reductions in mRNA levels from several PGC-1α neuron-specific target genes. The authors concluded that PGC-1α has a role in maintenance of gene expression programs for synchronous neurotransmitter release, structure, and metabolism in excitatory neurons (McMeekin et al., 2020). This would be consistent with our data in the cerebellum, in which granule cells are the most abundant glutamatergic neurons. It could also help explain the somewhat puzzling results that PGC1α deletion in parvalbumin-expressing inhibitory interneurons using a conditional deletion of exon 3 does not produce motor deficits (Lucas et al., 2014a). Combined with our data, this suggests that the SSR-SINE-exon2 isoform may normally act to co-repress genes in excitatory neurons, in this case glutamatergic neurons, to support motor behaviour. Nevertheless, how this isoform of PGC1α could repress, instead of co-activate, gene transcription remains unknown. The protein expressed from the SSR-SINE-exon2 isoform adds 6 amino acids to the N-terminus of PGC1α and replaces the 16 residues encoded by exon 1. Prediction of secondary structure reveals that this small amino acid change alters the protein structurally in that two alpha helices are replaced by a single larger alpha helix at the N-terminus of the protein (Fig. S5). It is possible that these changes alter the ability of this isoform to dock into NRs in such a way that prevents the recruitment of histone acetyltransferases, which would effectively inhibit transcription. Alternatively, it could be that the altered N-terminus of the protein recruits a histone deacetylase, or functions to repress activation of (yet unidentified) TFs in neurons by sequestering their partners, similar to how it alters NFkB signalling (Fig. 4F). Finally, another possibility is the SSR-SINE-exon2 isoform activates a protein that is a transcriptional repressor for the genes we find upregulated. More work is clearly required to understand how this novel brain-specific isoform can ultimately lead to the co-repression of genes and the extent to which this leads to alternations in neurotransmission.

Lastly, HIF1α was recently shown to activate transcription from the human B1 promoter, which corresponds to the SSR promoter in the mouse, by selectively interacting with it and not the reference promoter for the gene (Soyal et al., 2020). While the role of HIF1α in the brain has previously been limited to hypoxic insult, a recent study has shown that crucial polarity-controlled events in neuronal determination and cerebellar germinal zone exit – including spindle orientation during neural stem cell division, axon-dendrite specification, or adhesive events that promote synaptogenesis (Singh et al., 2016; Singh and Solecki, 2015; Uzquiano et al., 2018) are also regulated through HIF1α-dependent pathways, and, consequently, are sensitive to O_2_ tension (Kullmann et al., 2020). Whether brain-specific isoform of PGC1α that we identified participates in such events by interacting with HIF1α remains to be determined. As such, it will also be interesting to study whether the protein expressed from the SSR-SINE-exon2 isoform plays a role in the physiological outcomes associated with brain ischemia, including those related to stroke and pre-term birth.

In summary, we established that the novel brain-specific SSRSINE-exon2 containing isoform of *Pgc1α* is functional *in vivo* and that it has roles in brain physiology that differ from the reference isoform of the gene. The extent to which it influences neurological disease, how it interplays with the other isoforms of PGC1α in the different cell types in the brain and whether its interaction with the ER or PPARG contributes to sexual dimorphic phenotypes that influence psychiatric disease constitute promising areas for future experimentation.

## Materials and Methods

### SINE mutant animals

C57B6/J mice were purchased from The Jackson Laboratories. A single CAS9 target site (AATTGGAGCCCCATGGATGAAGG) was utilized to disrupt the ORF of the SINE-*Pgc1α* (SINE-Ppargc1a) variant. Complementary oligos were ordered from IDTDNA (Coralville, IA, USA) and cloned into a T7 sgRNA plasmid, and *in vitro* transcribed using Epicentre AmpliScribe T7 High Yield Transcription Kit (Madison, WI, USA). C57BL/6J one-cell embryos were microinjected with CAS9 SgRNA (10 ng/ul each) and 5’ capped/polyA tailed Cas9 RNA (100 ng/ul) derived from pCAG-T3-hCAS-pA, a gift from Wataru Fujii & Kunihiko Naito (Fujii et al., 2013). Microinjected embryos were surgically transferred to SWISS pseudo-pregnant females. At weaning, potential founders were screened by PCR amplicon sequencing (FWD: 5’-TGAGAATATCAGTCTCTGGGGGA-3’; Rev 5’-CAGCCCCTCCTCTGAAATACAAA-3’). Based on computationally predicted CAS9 off-target sites, the nearest genetically linked off-target site was nearly 27 mb away and contained 4 mismatches to the CAS9 target sequence, and therefore was not screened in the founder mice. Founders of interest were bred to wildtype C57BL/6 mice and F1 offspring were re-screened to confirm germline transmission. The mutant mouse line was crossed to wildtype C57BL/6J mice for at least two generations to eliminate any unknown, unlinked mutations. Phenotyping was done with founder line 4, which has a 4 bp deletion (TGAA) just 3’ of the SINE variant translational start site corresponding to chr5:51,912,715-51,912,718 (GRCm38/mm10 assembly). Mouse colony genotyping was done by primer/probe assay by Transnetyx (FWD-Primer 5’-AGGTTTTTTGCGAAAATCAGTGAACTAAT-3’; REV-Primer 5’-GCAGTTTGGAGCAATAGAGAAGAAC-3’; WT-PROBE 5’-AAAGTACCCTTCATCCATG-3’; Mutant-PROBE 5’-ACTTACAAAGTACCCTCCATG-3’). All animal protocols were approved by the Animal Care and Use Committee (ACUC) at the National Institute of Environmental Health Sciences (NIEHS) and experiments conducted in accordance with relevant guidelines and regulations. Female and male mice were included in all experiments, which were performed on age-matched WT and SINE homozygous littermates.

### PacBio sequencing and data analysis

We used RACE (Rapid Amplification of cDNA Ends) and whole brain RNA from the mouse to generate material for PacBio sequencing, which was performed at the National Institute Sequencing Core (NISC) in Bethesda. After sequencing, quality control was performed with FastQC (Available online at https://www.bioinformatics.babraham.ac.uk/projects/fastqc/). Primers and the first 50 bases with non-uniform nucleotide composition were removed with Trimmomatic (Bolger et al., 2014). A Phred quality filter was also applied, keeping sequences with quality over 10 (Q ≥ 10). To identify *Pgc1α* transcript variants, we artificially generated template sequences containing all possible exon combinations (198) and aligned the PacBio reads to this reference sequence using Minimap2 tool (Li, 2018). Only alignments with map quality over 20 (MAPQ ≥ 20) that aligned through the junction points of the template sequences (Exon-Exon, SINE-Exon and SSR-SINE-Exon) were kept. The alignments were visually inspected using IGV (Integrative Genomics Viewer) (Thorvaldsdottir et al., 2013).

### Antibody generation and specificity test

Antibodies were generated by Covance against the following amino acids of PGC1α: canonical isoform SQDSVWSDIEC, epitope MDEGYF within the SINE and NYGSSWETPSNQC within the SSR and those at position 513-526 at the C-terminus of the protein. Serum was utilized for the experiments shown herein in a dilution of 1-10. Specificity of antibodies was judged with purified peptides and in NIH3T3 cells expressing HA-tagged recombinant *Pgc1α*, which was cloned using forward 5’TTGACTGGCGTCATTCGGGA3’ and reverse 5’TCAGGAAGATCTGGGCAAAGAG3’ primers and expressed through the pInducer20 lentiviral vector (Addgene). Western blots were performed using actin as loading control; secondary antibodies were obtained from LiCOR and membranes were visualized using a LiCOR Odyssey imager.

### Histological analysis

Adult male and female SINE KO and WT littermate control mice were anesthetized with pentobarbital sodium and fixed via transcardial perfusion with 0.1M phosphate-buffered saline (PBS) followed by 4% paraformaldehyde (PFA) in PBS. Brains were removed, rinsed in PBS and fixed overnight in 4% PFA at 4°C. Brains were rinsed in PBS prior to cryoprotection in 30% sucrose in PBS. Cryoprotected brains were embedded in tissue-freezing medium (Triangle Biomedical Sciences) and sectioned. For each genotype and sex n=4 brains were sectioned at 25 µm in the sagittal or coronal plane and collected on Superfrost Plus microscope slides (Thermo Scientific, Waltham, MA). One set of slides was processed for 0.1% cresyl violet Nissl stain and a second with Luxol fast blue myelin stain. Following dehydration and clearing, slides were coverslipped with Permount (SP15-500, ThermoFisher Scientific) mounting medium.

### Behavioural tests

For the behavioural battery, animals generated at NIEHS were shipped to the UNC Mouse Behavioural Phenotyping Laboratory where 15 male and 11 female WT controls and 13 male and 6 female *Pgc1α* SINE isoform knockout (KO) were tested. Mice were 14 weeks in age at the start of behavioural testing. All animal care and procedures were conducted in strict compliance with the animal welfare policies set by the National Institutes of Health and by the University of North Carolina at Chapel Hill (UNC), and were approved by the UNC Institutional Animal Care and Use Committee.

### Social approach in a 3-chamber choice task

Mice were evaluated for the effects of *Pcg1*α SSR-SINE isoform deficiency on social preference. The procedure consisted of 3 10-minute phases: a habituation period, a test for sociability, and a test for social novelty preference. For the sociability assay, mice were given a choice between proximity to an unfamiliar, sex-matched C57BL/6J adult mouse (“stranger 1”), versus being alone. In the social novelty phase, mice were given a choice between the already-investigated stranger 1, versus a new unfamiliar mouse (“stranger 2”). The social testing apparatus was a rectangular, 3-chambered box fabricated from clear Plexiglas. Dividing walls had doorways allowing access into each chamber. An automated image tracking system (Noldus Ethovision) provided measures of time in spent in each chamber and entries into each side of the social test box. At the start of the test, the mouse was placed in the middle chamber and allowed to explore for 10 minutes, with the doorways into the 2 side chambers open. After the habituation period, the test mouse was enclosed in the center compartment of the social test box, and stranger 1 was placed in one of the side chambers. The stranger mouse was enclosed in a small Plexiglas cage drilled with holes, which allowed nose contact. An identical empty Plexiglas cage was placed in the opposite side of the chamber. Following placement of the stranger and the empty cage, the doors were re-opened, and the Behaviour in SSR-SINE KO mice 7 subject was allowed to explore the social test box for a 10-min session. At the end of the sociability phase, stranger 2 was placed in the empty Plexiglas container, and the test mouse was given an additional 10 min to explore the social test box.

### Morris water maze

The water maze was used to assess spatial and reversal learning, swimming ability, and vision. The water maze consisted of a large circular pool (diameter = 122 cm) partially filled with water (45 cm deep, 24-26° C), located in a room with numerous visual cues. The procedure involved two phases: a visible platform test and acquisition of spatial learning in the hidden platform task.

### Visible platform test

Each mouse was given 4 trials per day, across 2 days, to swim to an escape platform cued by a patterned cylinder extending above the surface of the water. For each trial, the mouse was placed in the pool at 1 of 4 possible locations (randomly ordered), and then given 60 sec to find the visible platform. If the mouse found the platform, the trial ended, and the animal was allowed to remain 10 sec on the platform before the next trial began. If the platform was not found, the mouse was placed on the platform for 10 sec, and then given the next trial. Measures were taken of latency to find the platform and swimming speed via an automated tracking system (Ethovision 15, Noldus, Wageningen, NL).

### Acquisition of spatial learning in the water maze via hidden platform task

Following the visible platform task, mice were tested for their ability to find a submerged, hidden escape platform (diameter = 12 cm). Each animal was given 4 trials per day, with 1 min per trial, to swim to the hidden platform. Criterion for learning was an average group latency of 15 sec or less to locate the platform, with a maximum of 9 days of training. Following testing on Day 9, mice were given a 1-min probe trial in the pool with the platform removed. Selective quadrant search was evaluated by measuring the number of swim path crosses over the previous platform location, versus the corresponding location in the opposite quadrant.

### Fear conditioning

Animals were held in an anteroom separated from the testing room to ensure that the animals did not hear testing of other animals for at least 30 min prior to training/testing. Training took place in four identical sound attenuating chambers (Context A; 28 x 21x 21 cm; Med-Associates Inc.). The floor of each chamber consisted of a stainless-steel shock grid (1/2 inch apart) wired to a shock generator and scrambler (Med-Associates Inc.) to deliver foot shocks. Mice were evaluated for learning and memory in a conditioned fear test (Near-Infrared image tracking system, MED Associates, Burlington, VT). The procedure had the following phases: training on Day 1, a test for context-dependent learning on Day 2, and a test for cue-dependent learning on Day 3. Two weeks following the first tests, mice were given second tests for retention of contextual and cue learning. Training. On Day 1, each mouse was placed in the test chamber, contained in a sound-attenuating box, and allowed to explore for 2 min. The mice were then exposed to a 30-sec tone (80 dB) that co-terminated with a 2-sec scrambled foot shock (0.4 mA). Mice received 2 additional shock-tone pairings, with 80 sec between each pairing.

### Context- and cue-dependent learning

On Day 2, mice were placed back into the original conditioning chamber for a test of contextual learning. Levels of freezing (immobility) were determined across a 5-min session. On Day 3, mice were evaluated for associative learning to the auditory cue in another 5-min session. The conditioning chambers were modified using a Plexiglas insert to change the wall and floor surface, and a novel odour (dilute vanilla flavouring) was added to the sound-attenuating box. Mice were placed in the modified chamber and allowed to explore. After 2 min, the acoustic stimulus (80 dB tone) was presented for a 3-min period. Levels of freezing before and during the stimulus were obtained by the image tracking system. Second test rounds were conducted 2 weeks after the first rounds.

### Accelerating rotarod, 5-min trials

At 16-19 weeks in age, subjects were given 2 tests for motor coordination and learning on an accelerating rotarod (Ugo Basile, Stoelting Co., Wood Dale, IL). The first test consisted of 3 trials, with 45 sec between each trial. Two additional trials were given 48 hours later. Retests were conducted when mice were 31-36 weeks and 50-61 weeks in age. Rpm (revolutions per minute) for each trial was set at an initial value of 3, with a progressive increase to a maximum of 30 rpm across 5 min (the maximum trial length). Measures were taken for latency to fall from the top of the rotating barrel.

### Rapid-reversal rotarod test, 5-min trials

At 56-66 weeks in age, subjects were given a 2-trial retest for motor coordination, using a rapid-reversal procedure. Rpm (revolutions per minute) for each trial was fixed at 10 rpm. Reversal of the direction of barrel spin occurred approximately every 15 s across the trial (maximum 5 min).

### Accelerating and Fixed speed Test on rotarod, 1 min trial

At 4-6 months of age in our initial cohort and at 32-37 weeks of age in the behavioural test battery cohort, mice were evaluated in a rotarod procedure, modified from a previously described protocol (Lucas et al., 2012). Mice underwent 2 trials each at rotating speeds of 0-10 (accelerating), 16, 24, 28, and 32 fixed rpm. Each trial was a maximum of 60 sec, with at least 5 min between each trial.

### Fixed speed test on the rotarod, 2-min trials

At 58-67 weeks in age the behavioural test battery cohort, mice were evaluated in a final rotarod procedure, modified from a previous ly described protocol (Lucas et al., 2012). Mice underwent 2 trials each at rotating speeds of 24, 28, and 32 fixed rpm. Each trial was a maximum of 120 sec, with at least 5 min between trials.

### Health status

Deficiency of the *Pgc1α* SINE isoform did not lead to overt changes in health or general motor ability. No subjects were lost from the behaviour study by the time of the final rotarod test, conducted when mice were age ∼60-70 weeks.

### Behavioural tests statistical analysis

For each procedure, measures were taken by an observer blind to mouse genotype. Behavioural data were analysed using one-way or repeated measures Analysis of Variance (ANOVA), with separate analyses for males and females. Fisher’s protected least-significant difference (PLSD) tests were used for comparing group means only when a significant F value was determined. For all comparisons, significance was set at p<0.05. Data presented in figures and tables are means (± SEM).

### Sample processing and RNA extraction

Mice were sacrificed and their brains immediately removed. While fresh, extracted brains were manually cleaned of brain stem tissue remnants by gross dissection, and split into cerebellum and rest of brain (whole brain minus cerebellum). Each specimen was then stored in an individual 15-ml conical tube, placed on dry-ice to snap-freeze and archived at −80C. To prevent cross-contamination and tissue degradation due to delayed processing, mice were dissected one at a time with a single-use scalpel blade each. RNA was extracted from the cerebellum or rest of the brain using TRIzol and 50-100 mg tissue per individual specimen; aqueous phase was retrieved to purify total RNA (ethanol precipitation, per manufacturer’s guidelines). Aliquots with up to 1 µg total RNA were treated with DNAse I in solution (Invitrogen) before utilizing for RT-PCR or microarrays.

### Microarrays and data analysis

The Affymetrix Human Genome U133 Plus 2.0 GeneChip® arrays were used to profile gene expression. Samples were prepared as per manufacturer’s instructions. Arrays were scanned in an Affymetrix Scanner 3000 and data was obtained using the GeneChip® Command Console and Expression Console Software (AGCC; Version 3.2 and Expression Console; Version 1.2) using the MAS5 algorithm to generate CHP-extension files. Analysis of variance (ANOVA) was used to identify statistical differences between means of groups at α<0.05 level among HG-U133 Plus 2.0 probe sets. Experiments followed a 2^3^-full factorial design with N=2 replication level, thus N=8 per group (WT or mutant) with 2 independent specimens each consisting of combinations across 3 variables (sex × genotype × organ) with 2 levels each (M vs. F, WT vs. Mut, Cb vs. Rest-of-Br). All specimens were matched for litter set as much as possible (i.e. mouse origin’s litters, parental breeding pair, or birth date). Ingenuity Pathway Analysis (IPA) was used to analyse differences between transcriptional profiles of SINE-mutant vs. WT littermates based on differential expression readouts with SNR>1 (|log2FC|>0.286) among the curated list of 1,980 single-gene differential probe sets (male cerebellum: 457; female cerebellum: 1,738; male rest-of-brain: 78; female rest-of-brain: 141).

### Data accessibility

Genomics data for this publication have been deposited in the NCBI’s Gene Expression Omnibus (Barrett et al., 2011; Barrett et al., 2013; Edgar et al., 2002) and are accessible through GEO Series accession number GSE152224.

## Author Contributions

FX generated and genotyped the initial animal cohort, performed the in-house rotarod assay, did the genomics analysis using publicly available data and qRT-PCR. OAL maintained the animal cohort, performed and analysed microarrays and qRT-PCR experiments. TW helped with bioinformatics. DG performed Western blots. BH and GR carried out transcript isoform reconstruction and analyses from long-range PacBio sequencing data. PJ performed histology and pathological assessment on dissected brains. SWS, KDS and JDC designed and performed behavioural studies. RPW and JHS conceptualized and oversaw the study, JHS was the lead writer of the manuscript.

## Acknowledgments

We thank the staff at the Core Facilities at NIEHS and NIH (Epigenetics and Genomics), Dr Artiom Gruzdev for generating the CRISPR/Cas9 KO animals and the critical comments on the manuscript by Drs. Paul Wade, G. Jean Harry and Kenneth Korach (NIEHS).

## Funding

This work was supported in part by the Intramural Research Program at the National Institute of Environmental Health Sciences of the National Institutes of Health (RPW), a doctoral scholarship from the Chilean National Agency for Research and Development (ANID, N°21201090) to BH and by Fondecyt grant 11140869 to GR (Chile).

## Conflict of Interest

The authors declare that the research was conducted in the absence of any commercial or financial relationships that could be construed as a potential conflict of interest.

## Supplementary Tables

https://orio.niehs.nih.gov/ucscview/Santos/Pgc1a/Table_S1_PacBio-Pgc1a.xlsx

https://orio.niehs.nih.gov/ucscview/Santos/Pgc1a/Table_S2_-_differentially_expressed_genes.xlsx

https://orio.niehs.nih.gov/ucscview/Santos/Pgc1a/Table_S3_-_KEGG_analysis.xlsx

https://orio.niehs.nih.gov/ucscview/Santos/Pgc1a/Table_S4_-_KEGG_candidate_NRs.xlsx

https://orio.niehs.nih.gov/ucscview/Santos/Pgc1a/Table_S5_-_striatum_data.xlsx

## Notes

### Competing Interest Statement

The authors have declared no competing interest.

