## supplemental data for "Mutations on a novel brain-specific isoform of PGC1α leads to extensive upregulation of neurotransmitter-related genes and sexually dimorphic motor deficits in mice"

**Supplemental Figures Legends**

**Figure S1. The antibody against the C-terminus of PGC1α is specific** (A) Antibodies were generated against the SSR, SINE, exon1-2 junction and the C-terminus (amino acids 513-526) of PGC1α. Peptides utilized to generate the 4 different antibodies were spotted on each membrane strip and incubated with each individual antibody (serum) separately. (B) NIH3T3 cells were transfected with a doxycycline-inducible HA-tagged vector expressing the canonical isoform of *Pgc1*α. Quantitative RT-PCR was used to estimate the expression level of *Pgc1*α among different clones; data was normalized to GAPDH. (C) Clone 1 (from B) was then used to test specificity of the antibodies generated; only the antibody with high specificity (against the C-terminus) is shown; anti-HA and actin were used as controls.

**Figure S2. CRISPR/cas9-generated SINE mutant mouse.** (A) Schematic representation of the mutations introduced in the SINE exon of the novel SSR-SINE-exon2 isoform. Upper track shows the endogenous *Pgc1*α locus with alternative SSR/SINE transcriptional start exons (black) and canonical exons 1 and 2 (gray) with splicing from SSR to SINE 2 to the reference Exon 2. Expanded SINE locus (65 bp) with the CAS9 target sequence (highlighted), translational start codon (underlined), and 4 bp deletion (double underline, dark highlight), which disrupts the SINE- *Pgc1*α variant ORF. Screening primers (FWD and REV) amplify a 670 bp fragment flanking the CAS9 target sequence. Bottom part shows the heteroduplex sequencing of the founder allele. (B) Quantitative RT-PCR using primers that span the junction between the SINE and exon 2 (left graph) were used to determine expression level of this isoform in whole brain of WT and mutant animals; N=3/group; each reaction was done in triplicates. Error bars depict ±SEM. GAPDH was used to normalize data; insert show representative qRT-PCR products ran on an agarose gel. Right graph shows data reflecting expression of the canonical (primers spanning exons 1-2) or total (primers spanning exons 2-3, common to all isoforms) levels of *Pgc1α* in the same animals. (C) Antibody against the C-terminus of PGC1α was used to probe its levels in the brain of WT and mutant animals; data are representative. NIH3T3 overexpressing recombinant HA-tagged PGC1α was used as positive control (+C).

**Figure S3. SINE mutant mice perform like WT littermates in most behavioural tests.** Animals were subjected to a battery of assays; unless otherwise stated data are means (± SEM). Open field: locomotor activity, vertical rearing, and time spent in the center of an open field was measured in a one-hour test. Sociability was measured in a 3-chamber choice for 10-min tests. *$p<0.05$, within-genotype comparisons. Contextual learning in a conditioned fear test. A) Mice were evaluated in a 5-min test 24hr following training. B) The second context test was conducted 2 weeks after the first test. Cue-dependent learning in a conditioned fear test. Arrows indicate onset of 3-min tone (80 dB). Mice were evaluated in a 5-min test for cue learning (Test 1) 48 hr after training. Levels of freezing were measured during the tone, presented 2 min after mice were placed in the modified conditioned fear chambers. Test 2 was conducted 2 weeks after the first test. Latencies to escape in the Morris water maze; means of 4 trials per day during a hidden platform test are depicted. Criterion for learning was a group average of 15 sec to find the escape platform. Only one group reached the learning criterion: female KO mice, on day 7 of testing.

**Figure S4. Gene expression analysis in the brains of WT and functional KO animals.** Overlap of probes differentially enriched in the cerebellum or rest of the brain (cerebrum minus cerebellum) between mutant and WT littermates based on gender. Red – female cerebellum, green – male cerebellum, blue – female rest of the brain and orange – male rest of the brain. N=4 per group.

**Figure S5. Secondary structures of the proteins derived from the reference or SINE-containing isoform of the *Pgc1α* gene are different.** Jpred4 (http://www.compbio.dundee.ac.uk/jpred4/) was used to predict secondary structures using the amino acid sequences of the reference PGC1α protein or that of the SINE-containing isoform. The red boxes represent α-helices while the green arrows are β sheets.

Fig S1

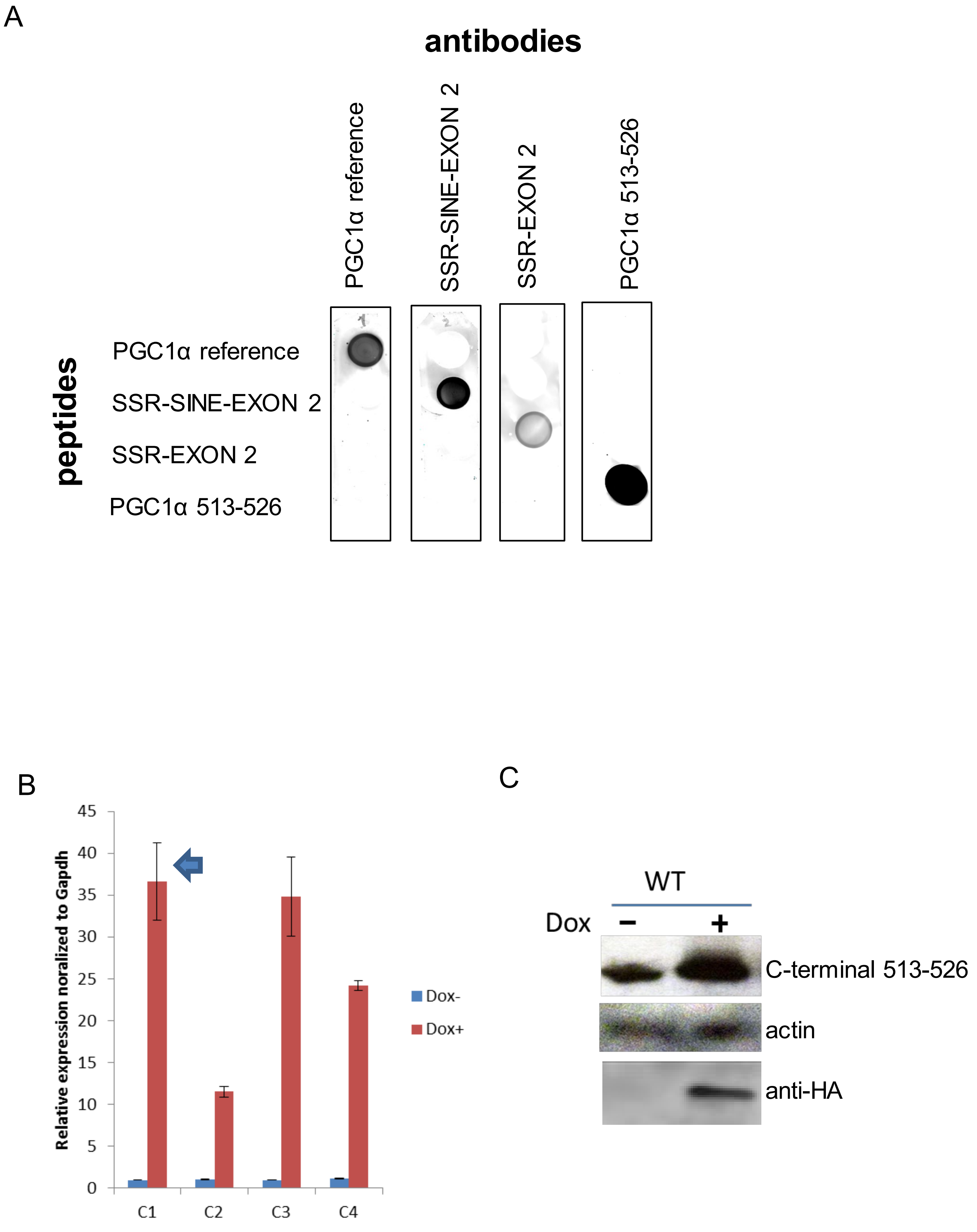

A
antibodies
PGC1α reference
SSR-SINE-EXON 2
SSR-EXON 2
PGC1α 513-526
peptides
PGC1α reference
SSR-SINE-EXON 2
SSR-EXON 2
PGC1α 513-526
B
Relative expression noralized to Gapdh
0
5
10
15
20
25
30
35
40
45
C1
C2
C3
C4
Dox-
Dox+
C
WT
Dox
−
+
C-terminal 513-526
actin
anti-HA

Fig S2

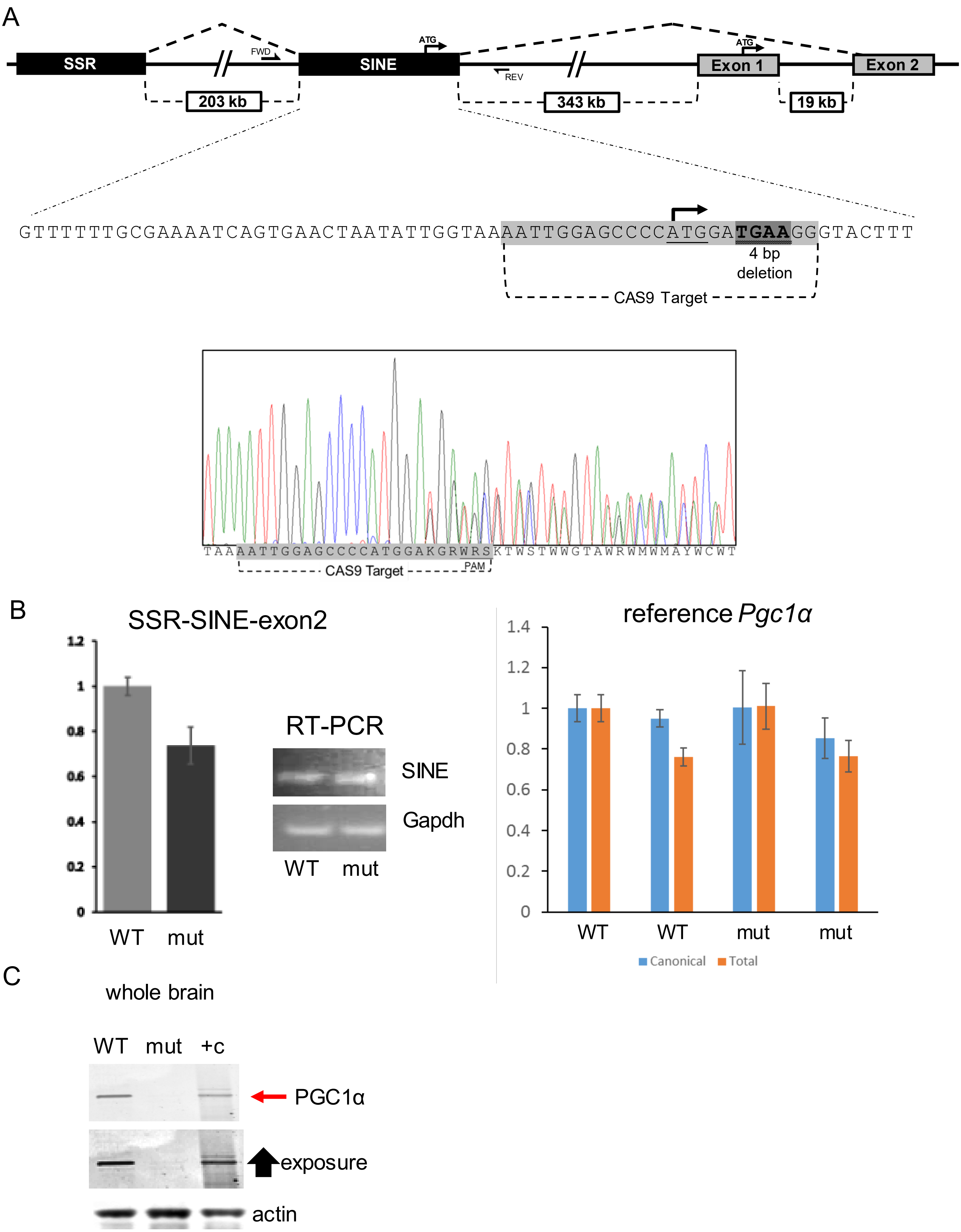
A
SSR
SINE
Exon 1
Exon 2
FWD
REV
ATG
ATG
203 kb
343 kb
19 kb
GTTTTTTGCGAAAATCAGTGAACTAATATTGGTAAAATTGGAGCCCCATGGATGAAGGGTACTTT
4 bp deletion
CAS9 Target
TAAAATTGGAGCCCCATGGAKGRWRSKTWSTWWGTAWRWMWMAYWCWT
CAS9 Target
PAM
B
SSR-SINE-exon2
WT
mut
RT-PCR
SINE
Gapdh
WT mut
reference Pgc1α
WT
WT
mut
mut
Canonical
Total
C
whole brain
WT mut +c
PGC1α
exposure
actin

Fig S3

### open field

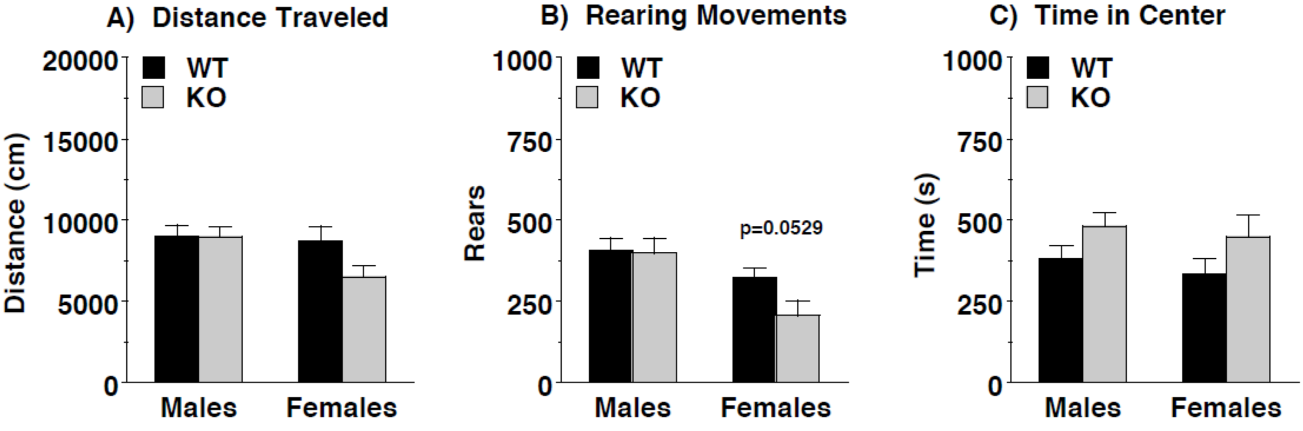


### Sociability three-chamber choice test

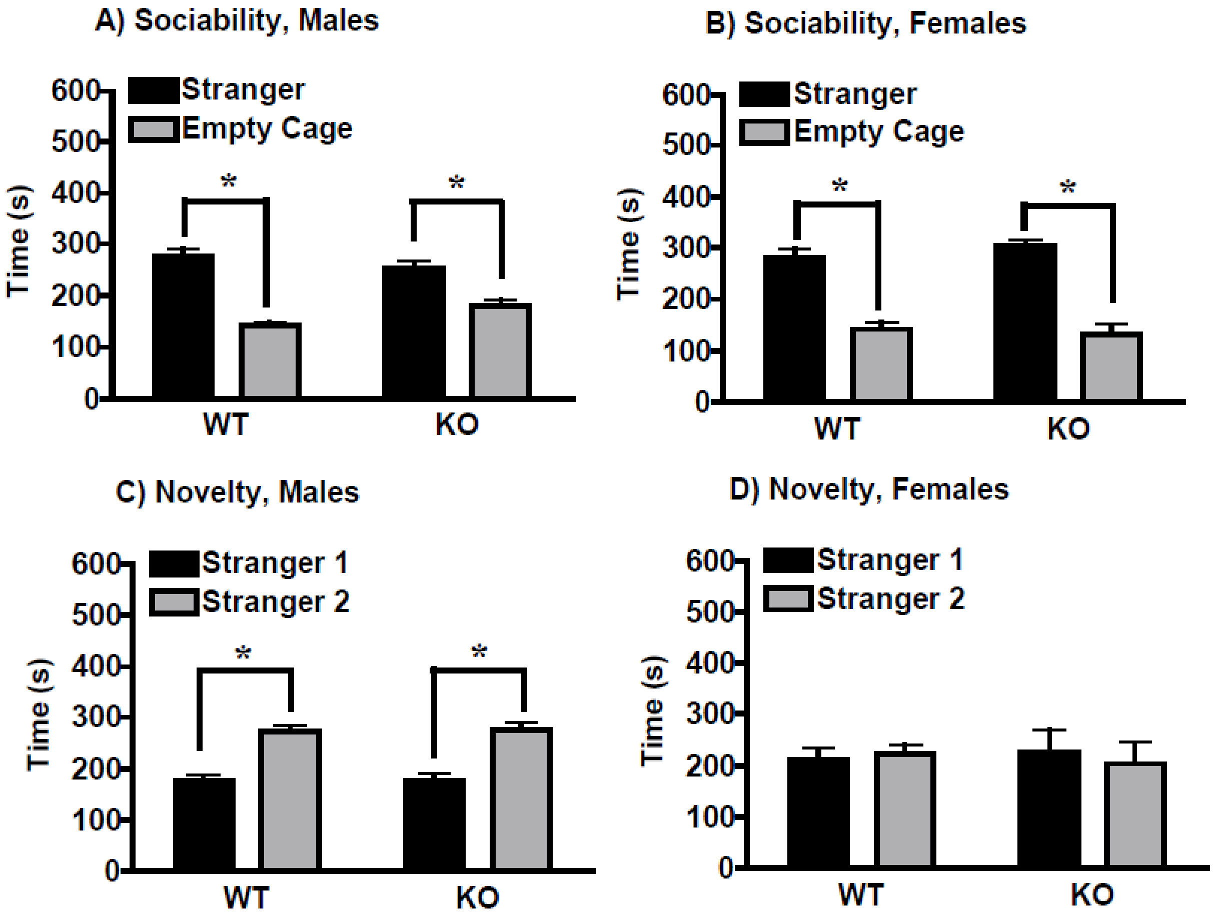

Fig S3

### Contextual learning – conditioned fear test

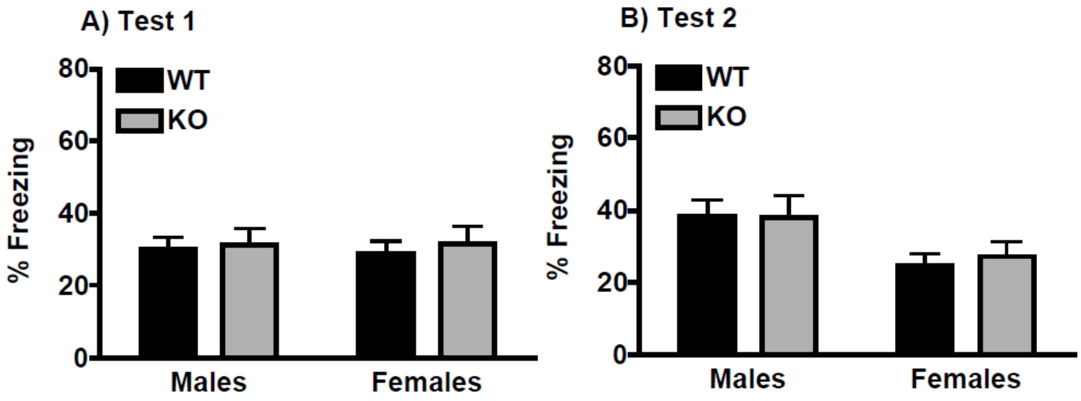


### Cue-dependent learning – conditioned fear test

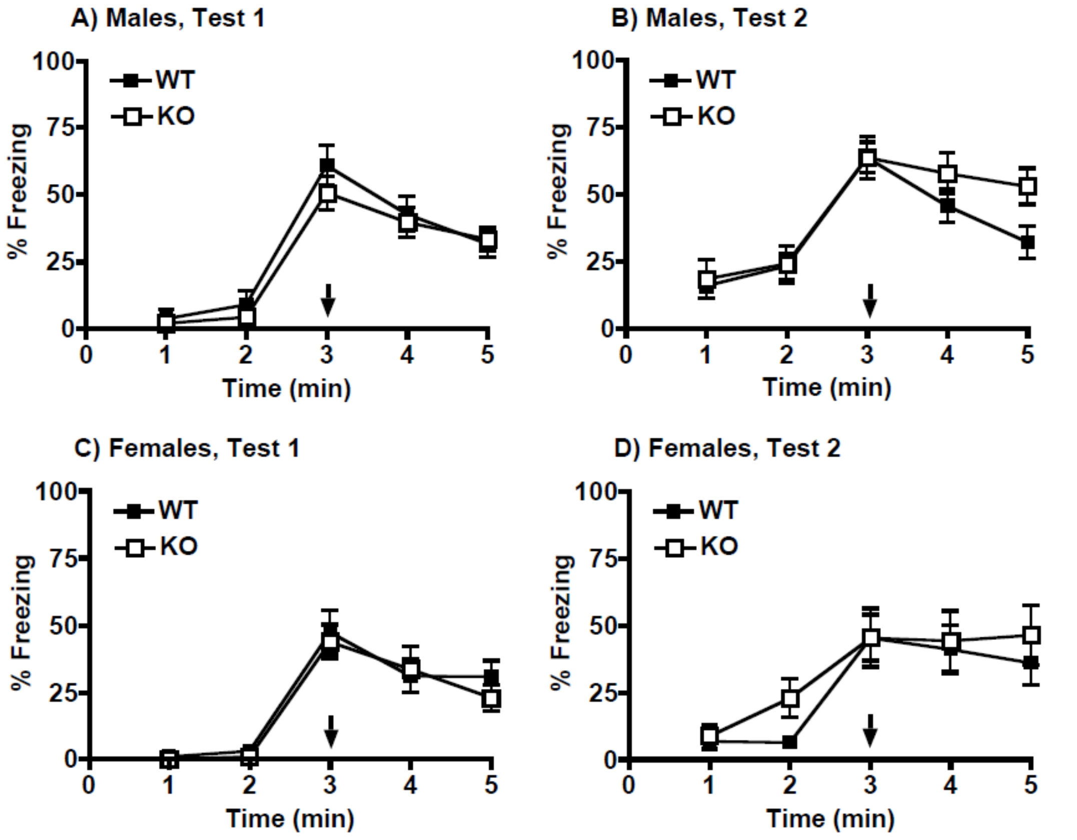

Fig S3

| | Males | | Females | |
|---|---|---|---|---|
| | WT | KO | WT | KO |
| Marble-bury assay | | | | |
| Number of marbles buried | 16 ± 0.4 | 16 ± 0.7 | 16 ± 0.8 | 15 ± 0.3 |
| Olfactory test | | | | |
| Latency to find buried food (s) | 74 ± 14[a] | 108 ± 19 | 117 ± 19 | 133 ± 46 |
| Hot-plate assay | | | | |
| Latency to respond (s) | 30 ± 0.3 | 28 ± 1.1 | 27 ± 1.1 | 30 ± 0.0 |

*p<0.05. [a]Data were removed for one male WT mouse that did not locate the buried food.

### Latency to escape and swim speed – water maze

| | Males | | Females | |
|---|---|---|---|---|
| | WT | KO | WT | KO |
| Visible platform, escape latency (s) | | | | |
| Day 1 | 25 ± 2 | 30 ± 2 | 30 ± 4 | 38 ± 7 |
| Day 2 | 14 ± 2 | 13 ± 2 | 19 ± 3 | 25 ± 6 |
| Swim speed (cm/s) | | | | |
| Day 1 of visible platform test | 17 ± 0.7 | 15 ± 0.6 | 17 ± 0.7 | 15 ± 1.0 |
| Day 1 of acquisition | 19 ± 0.4 | 16 ± 1.0* | 18 ± 0.7 | 16 ± 0.7 |

*p<0.05.

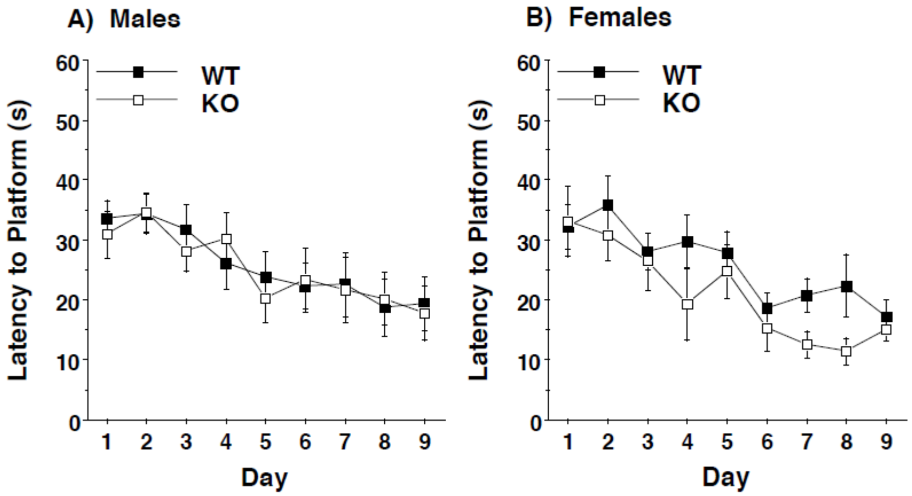

Fig S4

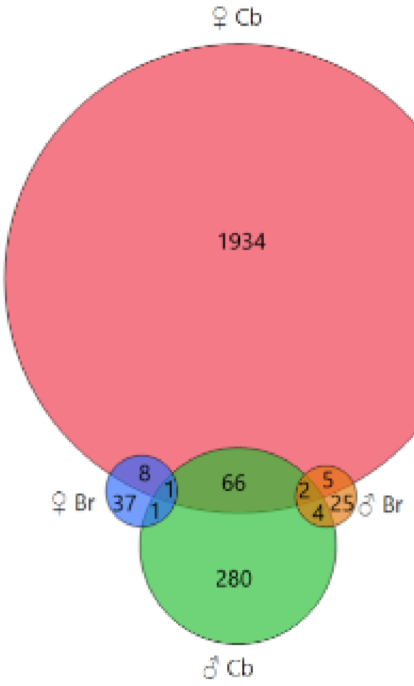

Fig S5

### Reference PGC1a – predicted secondary structure

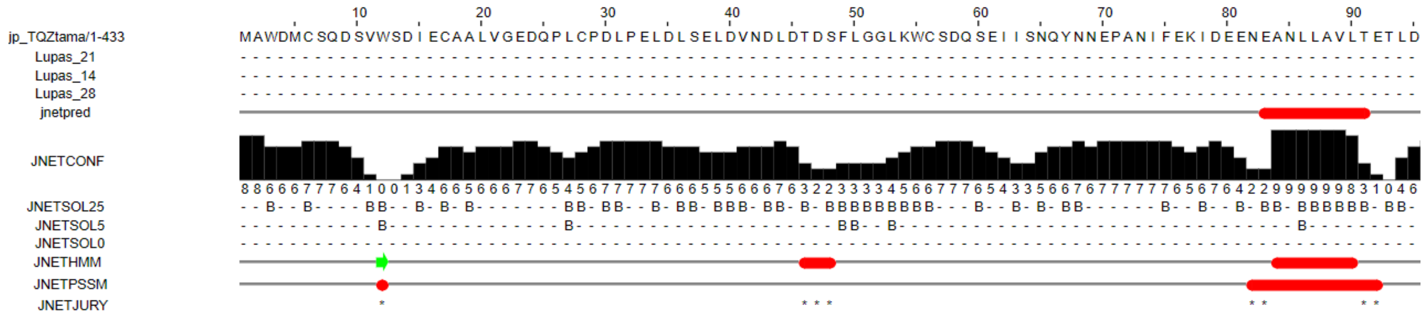


### SINE-PGC1a – predicted secondary structure

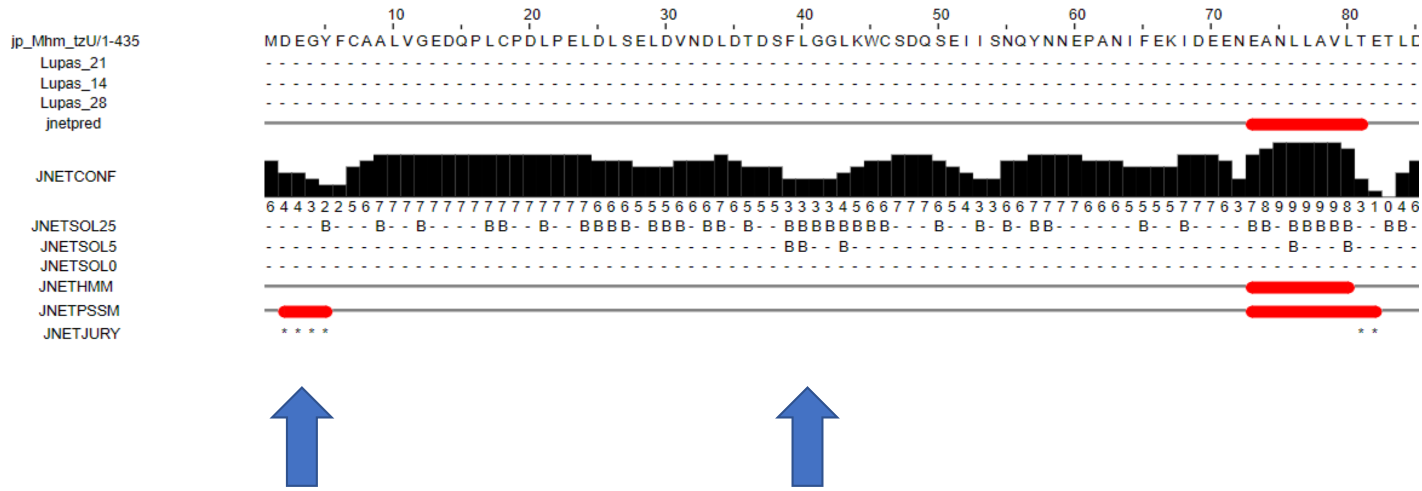
